# Protocol-dependent differences in IC_50_ values measured in hERG assays occur in a predictable way and can be used to quantify state preference of drug binding

**DOI:** 10.1101/534867

**Authors:** William Lee, Monique J. Windley, Matthew D. Perry, Jamie I. Vandenberg, Adam P. Hill

**Affiliations:** Victor Chang Cardiac Research Institute, 405 Liverpool Street, Darlinghurst, NSW 2010, Australia (WL, MW, MP, JV, AH); St Vincent’s Clinical School, University of NSW, Victoria Street, Darlinghurst, NSW 2010, Australia (WL, MW, MP, JV, AH)

## Abstract

Current guidelines around preclinical screening for drug-induced arrhythmias require the measurement of the potency of block of Kv_11.1_ channels as a surrogate for risk. A shortcoming of this approach is that the measured IC_50_ of Kv_11.1_ block varies widely depending on the voltage protocol used in electrophysiological assays. In this study, we aimed to investigate the factors that that contribute to these differences and to identify whether it is possible to make predictions about protocol-dependent block that might facilitate comparison of potencies measured using different assays

Our data demonstrate that state preferential binding, together with drug binding kinetics and trapping, is an important determinant of the protocol-dependence of Kv_11.1_ block. We show for the first time that differences in IC_50_ measured between protocols occurs in a predictable way, such that machine learning algorithms trained using a selection of simple voltage protocols can indeed predict protocol-dependent potency. Furthermore, we also show that a drug’s preference for binding to the open versus the inactivated state of Kv_11.1_ can also be inferred from differences in IC_50_ measured between protocols.

Our work therefore identifies how state preferential drug binding is a major determinant of the protocol dependence of IC_50_ measured in preclinical Kv_11.1_ assays. It also provides a novel method for quantifying the state dependence of Kv_11.1_ drug binding that will facilitate the development of more complete models of drug binding to Kv_11.1_ and improve our understanding of proarrhythmic risk associated with compounds that block Kv_11.1_.

## Introduction

The Kv_11.1_ (or hERG: human ether-a-go-go related gene) potassium channel carries one of the main repolarizing currents that contribute to the cardiac action potential – the rapid component of the delayed rectifier (I_Kr_) (Perrin *et al.*, 2008). Drugs that block Kv_11.1_ are the most common cause of acquired long QT syndrome (aLQTS), where drug-induced prolongation of repolarization can result in the fatal polymorphic cardiac arrhythmia Torsade de Pointes (TdP). As a result of these proarrhythmic side effects, a range of structurally unrelated drugs, including antihistamines, antibiotics, and antipsychotics were withdrawn from the market. In 2005, safety guidelines were put in place that mandated screening for Kv_11.1_ block for all new chemical entities in order to identify potentially pro-arrhythmogenic drugs during early preclinical development (Food and Drug Administration, HHS, 2005). While these guidelines have been successful in preventing new proarrhythmic drugs unknowingly coming to market, this has come at the cost of an unnecessarily high attrition rate of drugs in development (Sager *et al.*, 2014), i.e. current safety guidelines are very sensitive but not very specific.

At the preclinical level, a safety margin is determined by comparing the electrophysiologically determined inhibitory concentration of Kv_11.1_ (IC_50_) to the maximum plasma concentration (C_max_) of the drug, where the closer these two values are the higher the risk (Redfern *et al.*, 2003). One criticism of current guidelines is that they do not specify what methods, including cell type, voltage protocol, and temperature, should be used to measure IC_50_. Several studies have demonstrated significant differences in measured IC_50_ due to variations in all these factors (Kirsch *et al.*, 2004; Yao *et al.*, 2005; Milnes *et al.*, 2010; Windley *et al.*, 2018); in some cases as much as a 60-fold difference in IC_50_ was observed with different voltage protocols (Potet *et al.*, 2001; Rezazadeh *et al.*, 2004). A recent publication from the Comprehensive *in vitro* Proarrhythmia assay (CiPA) initiative acknowledged this issue and noted that it is not possible to predict protocol dependence in advance (Fermini *et al.*, 2016). Therefore, if a ‘true’ IC_50_ value cannot be reliably determined, then a safety margin is not an effective way of identifying and eliminating risk (Lee *et al.*, 2017).

To address this issue, CiPA has been proposed as a new safety paradigm in understanding TdP and assessing proarrhythmia risk (Sager *et al.*, 2014). One of CiPA’s key objectives is to use detailed *in vitro* electrophysiological characterization of drug interaction with Kv_11.1_ to build *in silico* models to predict arrhythmia risk. One factor that is likely important for this approach, is understanding the state dependence of binding (Lee *et al.*, 2017). It has been shown that for some drugs, the affinity can be as much as 30-fold greater for the inactivated state (K_i_) relative to the open state (K_o_) (Suessbrich *et al.*, 1997; Ficker *et al.*, 1998; Perrin, Kuchel, *et al.*, 2008; Wu *et al.*, 2015) whereas other drugs have no apparent state preference (Hill *et al.*, 2014). Furthermore, previous studies have shown that two drugs with opposite state preference (10^4^ fold difference in open vs inactivated state affinity) but the same IC_50_ can cause up to 56ms (~15%) difference in the extent of prolongation of the action potential duration (APD) at an IC_50_ dose (Lee *et al.*, 2017). Thus, state preference is an important consideration when determining whether a Kv_11.1_-drug interaction is likely to be proarrhythmic.

In this study, we use both *in silico* and *in vitro* approaches to explore the extent to which protocol dependent differences in Kv_11.1_ drug binding affinity can be explained by state dependent drug binding. Additionally, we demonstrate that 1) measuring potency with a range of voltage protocols that result in different ratios of state occupancy can be used to predict and quantify a drug’s state dependent drug binding characteristics; and 2) That a drugs potency measured using specific individual protocols can be predicted from IC_50_s measured using other protocols. Importantly, these either of predictions can be made based on simple voltage protocols that are readily amenable to automated, high thoughout patch clamp platfomrs. Our work provides a novel method for quantifying the state dependence of Kv_11.1_ drug interaction from simple measures of potency and identifies how state preferential drug binding is a major determinant of the protocol dependence of IC_50_ measured in preclinical Kv_11.1_ assays.

## Materials and Methods

### In silico modelling

A Markov state model of drug binding to Kv_11.1_ (Figure 1A and Supplementary figure 1) was used to simulate drug block of I_Kr_ (Lee *et al.*, 2016). This model has 3 closed states (C_0_,C_1_,C_2_), an open-conducting state (O) and an inactivated state(I), and drugs can bind to either the open state (OD) and/or the inactivated state (ID). Transitions between the non-drug bound states are expressed as rate constants (*k*_f_ for the forward rate and *k*_b_ for the backward rate) and are of the format shown in equations 1 and 2:

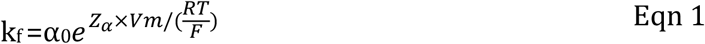

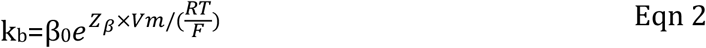

where R is the gas constant 8.314, T is the temperature in Kelvin, F is Faraday’s constant, V_m_ is the membrane voltage in millivolts. α_0_, *Z*α, β_0_, *Zβ* for the model at 37 °C are shown in Supplementary Figure 1. To approximate gating at 22 °C (Figure 9), individual rates (*k*_f_ or *k*_b_) were scaled for temperature using to Q_10_ values of 2.1, 1.7, 2.5 and 2.6 for activation, deactivation, inactivation and recovery from inactivation transitions respectively as previously reported (Vandenberg *et al.*, 2006).

**Figure 1.**
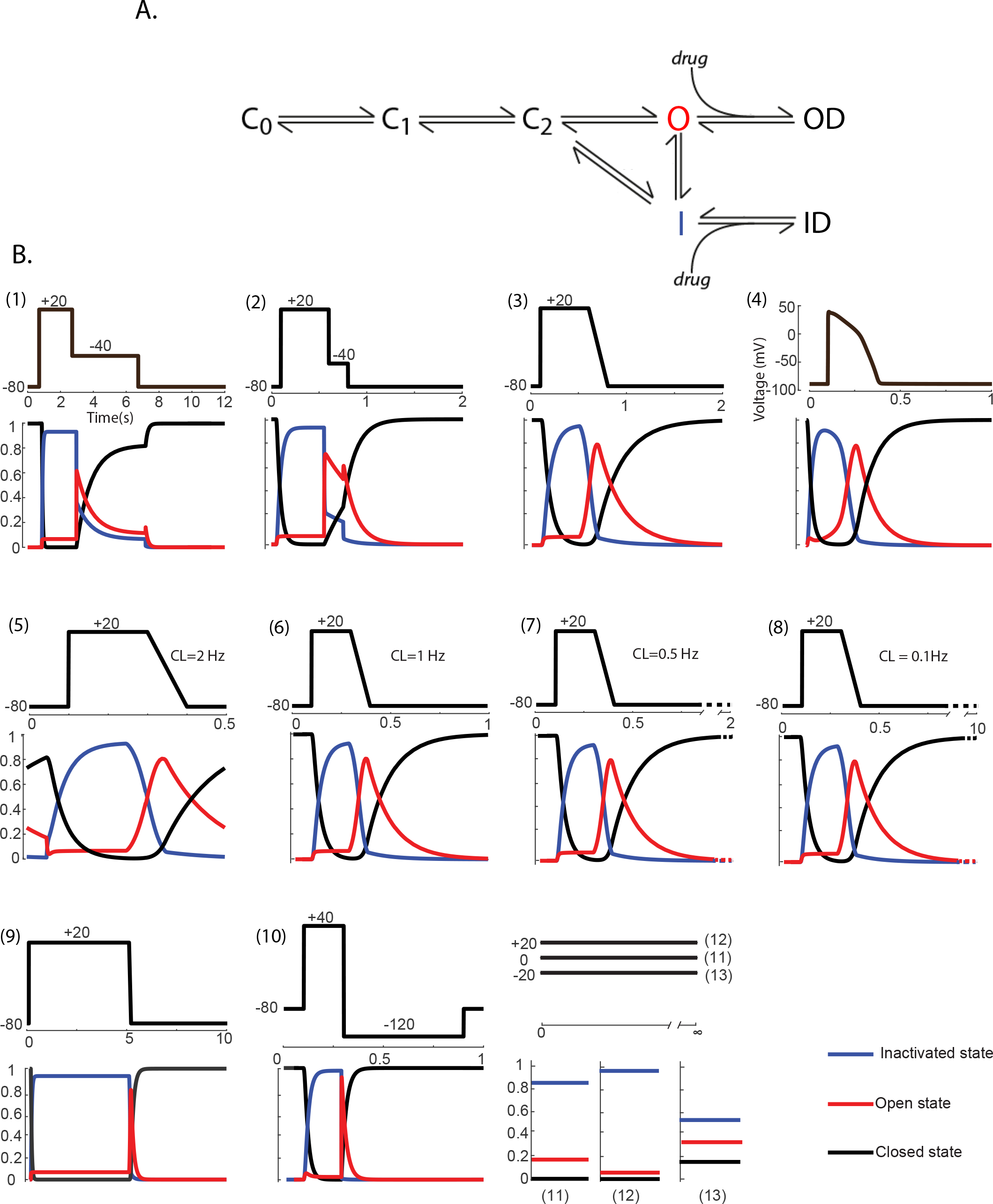
A) Markov state model of Kv_11.1_-drug interaction. Parameters describing voltage-dependent Kv_11.1_ gating transitions are presented in Supplement Figure 1. (B) Voltage waveforms and their gating state occupancies (open (red), inacticated (blue), closed (black)) for each of the 13 voltage protocols used.

Drug binding was described as forward and reverse rates for binding to the open state (k_f,open_ and k_b,open_ respectively) and to the inactivated state (k_f,inact_ and k_b,inact_ respectively). Drug association constants for the open state (K_o_) and the inactivated state (K_i_) were calculated as the ratio of the forward and reverse binding rates:

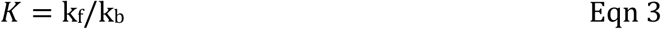

The relative drug affinity (K_O/I_) for the two states was described as a ratio of each individual state’s drug association constant as shown in equation 4. Where K_O/I_ <0.5 indicates a greater relative affinity for the inactivated state, K_O/I_ >2 indicates a greater relative affinity for the open state and 0.5 < K_O/I_ <2 indicates drugs that have equal affinity for both states or minimal difference in affinity for either state.

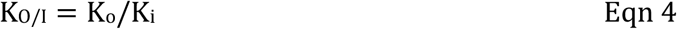

### Theoretical drugs

To examine the effect of state dependent drug binding on measured potency we used the same set of 6561 theoretical drugs specified in Lee et al. (Lee et al 2016). This was created through permutations of the 2 forward and 2 reverse rates for drug binding in the range of 0.01 – 100s^−1^ using half logarithmic increments (0.01, 0.03, 0.1, 0.3, 1, 3, 10, 30, 100s^−1^) for each drug state transition. For machine learning training and validation we used a separate set of 2000 and 1000 theoretical drugs respectively. In this case, forward and reverse rates of binding were randomly generated in the range of 10^4^ – 10^8^ s^−1^ for forward rates, and 0.001-100s^−1^ for reverse rates. These rates were consistent with previously published *in vitro* constrained kinetic data (Windley *et al.*, 2017) and produced IC_50_ values in a physiological range of the order 10^−10^ – 10^−3^ M.

Drug binding simulations were performed using the above described sets of theoretical drugs over a wide range of drug concentrations (10^−8^ to 10^7^M for the initial dataset, or 10^−15^ to 10M for the machine learning dataset). Current was normalized to the control current. For pulsed voltage protocols this was measured as peak current, while for non-pulsed protocols was taken as mean current over >100ms once the current had reached equilibrium. IC_50_ values were calculated from dose response curves using the Hill equation (equation 5):

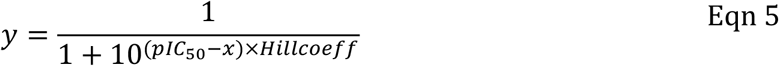

Where *y* is the normalized peak current amplitude, *x* is the log_10_ drug concentration in M, *pIC_50_* is the log_10_ IC_50_ in M and *Hillcoeff* is the hill slope of the dose response curve. Protocol to protocol differences in IC_50_ were measured as the Δlog[IC_50_], which is the difference in log_10_ [IC_50_] for each protocol. All simulations and analyses were performed using Matlab software (Mathworks, Natick, MA).

### Cell Culture

Chinese hamster ovary (CHO) cells stably expressing Kv_11.1_ (cell line PTA-6812) were purchased from the American Type Culture Collection (Manassas, VA). Cells were cultured in Hams F12 nutrient mix (Thermo Fisher Scientific, Waltham, MA) with added 5% fetal bovine serum (Sigma-Aldrich, Sydney, Australia) and maintained at 37°C with 5% CO_2_.

### Patch clamp Electrophysiology

Kv_11.1_ currents were recorded in the whole-cell voltage-clamp configuration at 22°C. The current signal was sampled at 5kHz and filtered at 1kHz with an Axopatch 200B amplifier (Molecular Devices, Sunnyvale, CA) interfaced with a Digidata 1440A analog-to-digital convertor (Molecular Devices, Sunnyvale, CA). Series resistance was compensated by at least 80% in all experiments. External bath solution contained (in mM): 130 NaCl, 5 KCl, 1 MgCl_2_, 1 CaCl_2_, 12.5 glucose, 10 HEPES adjusted to pH 7.4 with NaOH. Single-use patch pipettes were made from borosilicate glass (Havard Apparatus, Holliston, MA) and pulled using a vertical two-stage puller (PP-830, Narishige, Tokyo, Japan) with resistance of 2-4MΩ. Pippettes were filled with internal solution containing (in mM): 120 potassium gluconate, 20 KCl, 1.5 Mg_2_ATP, 5 EGTA and 10 HEPES, adjusted to pH 7.4 with KOH. Data was corrected for a calculated liquid junction potential of −15mV. Data was acquired with pCLAMP 10 acquisition software (Molecular Devices, Sunnyvale, CA), Leak currents were subtracted offline and data analysis was performed using Clampfit (Molecular Devices, Sunnyvale, CA), Prism (V7; GraphPad, San Diego, CA), and Matlab (Mathworks, Natick, MA).

### Pharmacology

Verapamil, terfenadine, cisapride and clozapine were purchased from Sigma-Aldrich, (Sydney, Australia) and dissolved in dimethylsulfoxide (DMSO). The maximum final concentration of DMSO in the recording solution was less than 0.01% (v/v), which is well below the 0.1%(v/v) shown to have no effect on Kv_11.1_ channel activity (Walker *et al.*, 1999). Drugs were delivered via a Dynaflow Resolve microfluidic device (Cellectricon, Mölndal, Sweden) allowing delivery of discrete solutions of various drug concentrations under laminar flow with a solution exchange time of less than 30 ms. Current amplitudes were measured when steady state had been achieved. Five drug concentrations were used to construct a Hill curve and calculate an IC_50_ value for verapamil, cisapride and clozapine. Three concentrations were used for terfenadine as drug binding has very slow kinetics and we wished to avoid significant current run-down whilst allowing sufficient time for drug binding to reach steady-state.

### Voltage Protocols

We studied 13 voltage protocols that reflected the range of commonly used published voltage protocols as well as incorporating variations in cycle length, which allowed sampling of the Kv_11.1_ channel at different amplitudes and durations of state occupancy. Voltage waveforms and their corresponding state occupancies for the open (red), inactivated (blue) and closed (black) states are shown in Figure 1B. All protocols had a holding potential (V_h_) of −80mV unless otherwise stated. Protocol 1 was a 2 step protocol with an initial step (V_1_) to +20mV for 2s followed by a −40mV step (V_2_) for 4s and a total interpulse time (CL) of 12s. (Milnes *et al.*, 2010). Protocol 2 was also a 2 step protocol with V_1_= +20mV for 500ms, V_2_=−50mV for 200ms and CL=2s (Yao *et al.*, 2005). Protocol 3 was step-ramp protocol with a V_1_= +20mV for 500ms, followed by a repolarizing ramp (V_r_) of −0.5V/s, CL=2s (Yao *et al.*, 2005). Protocol 4 was the voltage waveform of an O’Hara Rudy action potential (O’hara *et al.*, 2011). Protocols 5-8 consisted of a step-ramp with V_1_=+20mV for 200ms, V_r_=1V/s and CL of 500ms, 1s 2s and 10s respectively. Protocol 9 was a step-ramp protocol with a long V_1_ step of 5s at +20mV, followed by V_r_ =1 V/s and CL=10s. Protocol 10 was a 2 step protocol with V_1_=+40mV for 200ms, V_2_ =−120mV for 600ms and CL=1s, resulting in a negative hook-tail current (Perrin, Kuchel, *et al.*, 2008). Finally Protocols 11-13 were performed with direct application of the drug during voltage holding at V_h_= 0mV (Hill *et al.*, 2014), +20mV and −20mV respectively, CL=∞. For *in vitro* experiments and *in silico* machine learning algorithms, a subset of 6 of the above protocols were used; this included Protocols: 1, 8, 10, 11, 12 and 13. Specific details of individual protocols as well as evoked Kv_11.1_ currents from *in silico* and *in vitro* experiments are shown in the supplementary data.

To quantify the state occupancy fraction of open (Fso_open_) and inactivated (Fso_inact_) states for each voltage protocol, a time-integral of state occupancy was calculated for each state as follows:

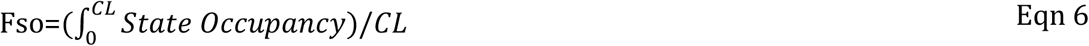

Where *State Occupancy* represents the state occupancy for each of the 3 states (Open=O, Inactivated=I, or Closed=C_1_+C_2_+C_3_) defined in the Markov Model above in the section titled *In silico modelling*, and CL is the total interpulse time as stated above. The ratio of open versus inactivated state occupancy (R_O/I_) for each voltage protocol was therefore defined in equation 7:

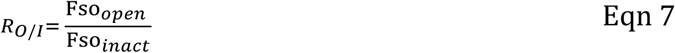

### Machine learning

A subset of 6 voltage protocols was chosen which maximized the differences in state occupancy for the three states and demonstrated the largest variation in *in silico* IC_50_ measurements. This subset was used to simulate and measure IC_50_‘s using the randomly generated 2000 training and 1000 validation theoretical drug sets. These sets of IC_50_ values were then used to train and validate machine-learning algorithms to predict drug binding characteristics including state preference and protocol dependent affinity. All machine learning prediction algorithms were generated and validated using Matlab Neural Network fitting application (Mathworks, Natick, MA). Training datasets were normalized (z-score) to have a mean of 0 and a standard deviation of 1. Validation data sets were z-score transformed using the mean and standard deviation of their respective training set. Neural networks were constructed using with between 10-100 neurons within one hidden layer. The neural network which best fitted the training data set was then tested for overfitting using the corresponding validation data set.

### Statistical Methods

Analysis of variance (ANOVA) was used for statistical analysis between normally distributed data for multiple groups. P values of <0.05 were considered significant. Statistical analyses were performed in Matlab (Mathworks, Matick, MA).

## Results

### In silico variations in measured drug affinity to Kv_11.1_

To test the influence of state preferential drug binding on variation in measured Kv_11.1_ affinity we simulated block of Kv_11.1_ using 2 voltage protocols that have been used in previously published studies (Milnes *et al.*, 2010; Hill *et al.*, 2014). We simulated dose response curves for an inactivated state preference drug with a 1000-fold higher affinity for the inactivated state vs open state (K_O/I_ =10^−3^). Example simulated currents in response to increasing concentrations of the inactivated state-preference drug for the two protocols are shown in Figure 2B. For each protocol a Hill curve was fit to the simulated data to derive an IC_50_ value for channel block. The inactivated state preference drug had a 50-fold higher affinity for Kv_11.1_ when measured using the +20mV V_hold_ protocol 12 compared to the 0.1Hz step-ramp protocol 8 (Figure 2C left panel; IC_50_=1mM and 50mM respectively. As a comparison we also simulated block using a non-state preference drug which showed no difference in Kv_11.1_ affinity between the two protocols (IC_50_=100mM) (Figure 2C right panel).

**Figure 2.**
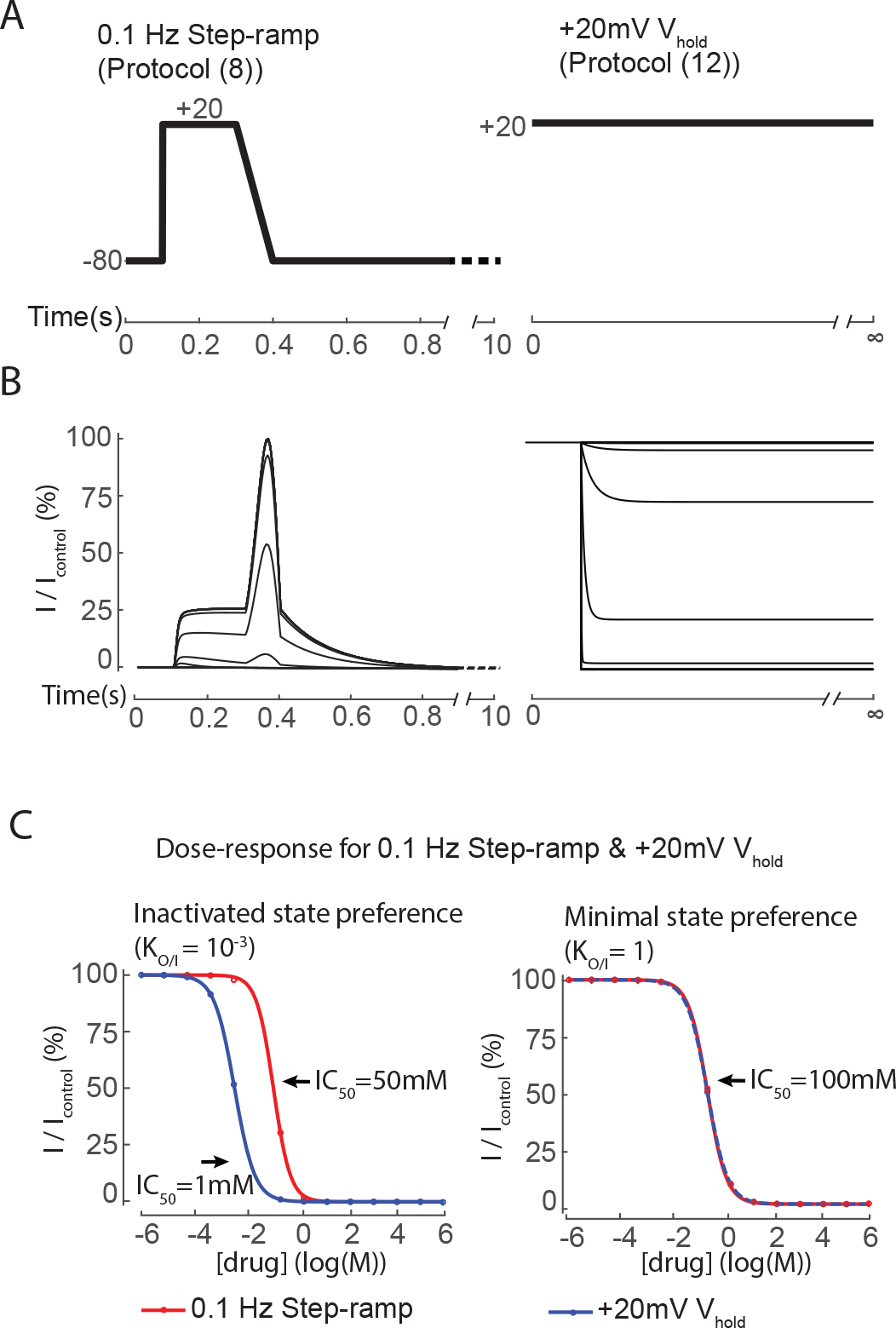
A) Voltage waveforms for the 0.1 Hz step-ramp protocol (8) and the non-pulsed +20mV V_hold_ protocol (12). B) Examples of *in silico* simulated Kv_11.1_ currents evoked when voltage clamped using voltage protocols 8 (left) and 12 (right), in the presence of increasing concentrations of drug (10^−6^ to 10^6^M is shown). C) Dose response curves derived from normalized peak current amplitude shown in panel B). The left side graph demonstrates a 50-fold difference in measured IC_50_ between the 2 protocols for a drug that preferentially binds to the inactivated state (K_O/I_ =10^−3^). In comparison, the right side graph shows a drug that has no state preference (K_O/I_ =1) has no difference in measured IC_50_ between the 2 protocols.

To probe the protocol dependence of measured Kv11.1 affinity in more detail, we extended our study to a panel of 6561 theoretical drugs with varying state preference and a larger array of 13 voltage protocols. For each combination of drug and protocol, a dose response curve was simulated (as shown in Figure 2B) and an IC_50_ derived through fit of the Hill equation. For each drug we then calculated the difference in log IC_50_ (ΔIC_50_) between pairs of protocols and grouped them with respect to state preference (where inactivated state preference: K_O/I_ <0.5 are colored blue, open state preference: K_O/I_ >2 are colored red, and minimal-state preference: 0.5< K_O/I_ <2 are colored yellow) (Figure 3). This pairwise analysis of protocols gave a total of 78 comparisons.

We first examined the distribution of ΔIC_50_ measured using the same two protocols shown in Figure 2 (protocols 8 and 12, Figure 3A). We observed a wide range of ΔIC_50_, from −0.9 to 1.7 which equates to a difference in IC_50_ for protocol 8 vs protocol 12 of 8-fold less or 50-fold higher. Overall, drugs tended to be grouped by state preference. While all drugs tended to have a higher affinity when measured with protocol 12, this effect was most pronounced for inactivated state preference drugs. For reference, the inactivated preference drug example from Figure 2 is marked as an (*) and the non-state preference drug is marked as a (†). Other protocol comparisons gave much clearer separation of drugs according to their state preference. A pairwise comparison of protocols 5 and 12 (Figure 3B), resulted in a clear bimodal separation of drugs based on open:inactivated state preference with inactivated state preference drugs displaying a higher affinity and open state preference drugs having a lower affinity for protocol 12 relative to protocol 5. Consistent with our previous observations, non-state preference drugs had minimal or no difference in measured IC_50_ between protocols. It is important to note that subtle differences in the protocols being compared can significantly affect the observed distributions for ΔIC_50_. For example, in Figure 3A and B, Protocol 12 is compared against protocols 8 and 5 respectively. The only difference between protocols 5 and 8 is the interpulse time (CL=500ms vs 10000ms), yet the observed distributions (compare Figure 3A and 3B) are clearly different, highlighting that it is not solely the ‘test pulse’ component of the protocol that is important in defining the measured potency. Conversely, when 5 and 8 are compared against each other (Figure 3C), there is no observable clustering of drugs based on their state preference. However, the fact that there were still observable differences in ΔIC_50_ between two protocols for different drugs (maximum ΔIC_50_ of −1.3 or +0.1), suggests that state preference of binding is not the only factor that influences protocol dependent affinity.

Finally, we examined an example of a pairwise comparison between non-pulse protocols. A comparison of ΔIC_50_ for protocols 12 and 13 – static holding potentials of +20mV and −20mV respectively - is shown in Figure 3D. We observed a distinct grouping of drugs based on state preference, albeit over a smaller range of ΔIC_50_; from −0.7 to 0.2 difference between the 2 protocols. In this case, inactivated state preference drugs had a higher affinity for protocol 12 and open state preference drugs for protocol 13. It is notable that these two protocols show the greatest separation of drugs based on state preference as they are pure equilibrium measures meaning there is no cycling of state occupancy between the open, inactivated and closed states during the protocol and consequently there is no influence from drug binding kinetics.

**Figure 3.**
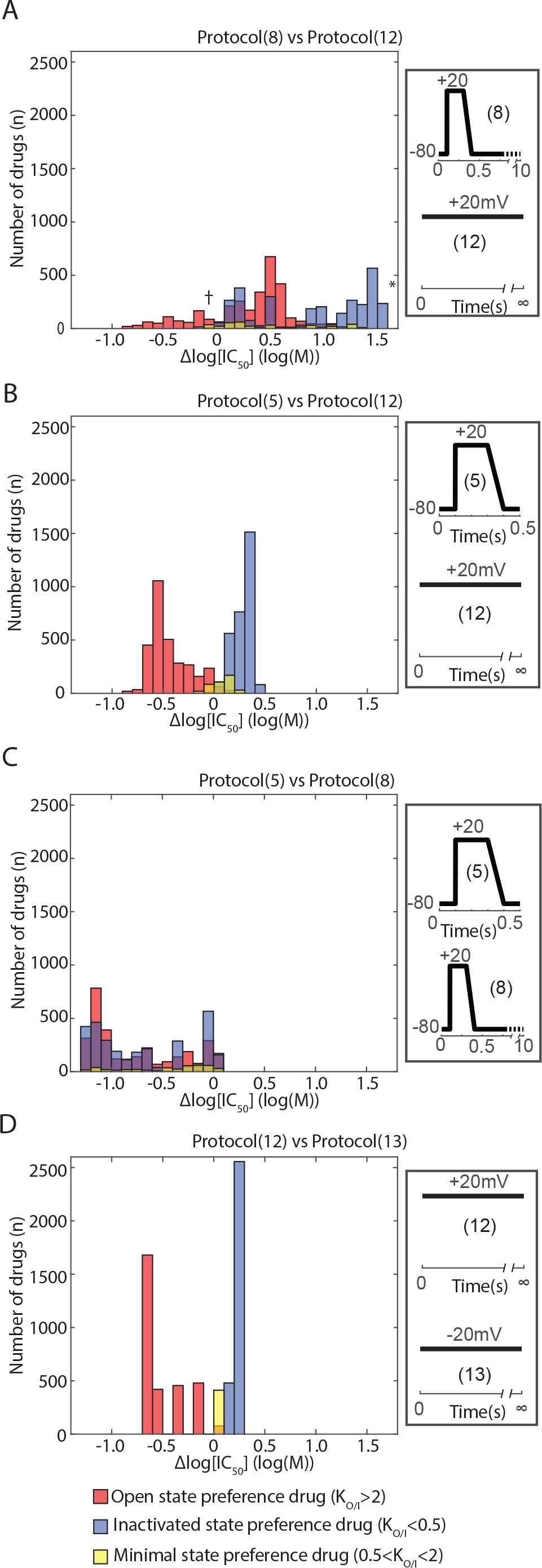
Differences in IC_50_ between 2 protocols for 6561 theoretical drugs. Drugs are grouped by preferential binding to the open state (K_O/I_ >1, red), inactivated state (K_O/I_ <1, blue) and minimal state preference (0.5< K_O/I_ <2, yellow). A) Shows a wider landscape of comparison between protocol (8) and protocol (12) from Figure 3. The inactivated state preference drug and non-state preference drug examples from Figure 3 are marked as a (*) and (†) respectively. The difference in IC_50_, measured as the difference in log_10_[IC_50_], (Δlog[IC_50_]) between these two protocols spans over a wide range with some grouping with respect to drug state preference. B) Comparison between IC_50_s from protocol (5) and protocol (12): showing more distinct grouping with respect to state preference. C) Comparison between IC_50_s from protocol (5) and protocol (8): no obvious grouping is seen with respect to drug state preference due to the very similar waveform of the 2 voltage protocols. But up to 20-fold variation in IC_50_ is still observed indicating mechanisms other than state-preference can influence drug affinity to Kv_11.1_. D) Comparison between IC_50_s from protocol (12) and protocol (13): very distinct grouping with respect to drug state preference is seen over a smaller range. The right-side panels (A-D) show the waveforms of the two voltage protocols being compared for their respective histogram.

Given this varied separation of drugs with different pairs of protocols, we next investigated which pairwise comparison gave the greatest ΔIC_50_ across all our drugs. To do this we took the maximum ΔIC_50_ from each pairwise comparison for each drug (Figure 4). This demonstrated that the maximum ΔIC_50_ was most often observed for comparisons between protocols 8 and 12, as well as protocols 8 and 13 (Figure 4A). More specifically, we observed that open state preference drugs tended to have a maximum ΔIC_50_ between protocols 8 and 13 whilst inactivated state and non-state preference drugs tended to have a max ΔIC_50_ between protocols 8 and 12 (Figure 4B). This supports the idea that individual voltage protocols sample distinct gating state ‘space’ and hence differentially interact with state preferential drug binding to determine these shifts in IC_50_.

**Figure 4.**
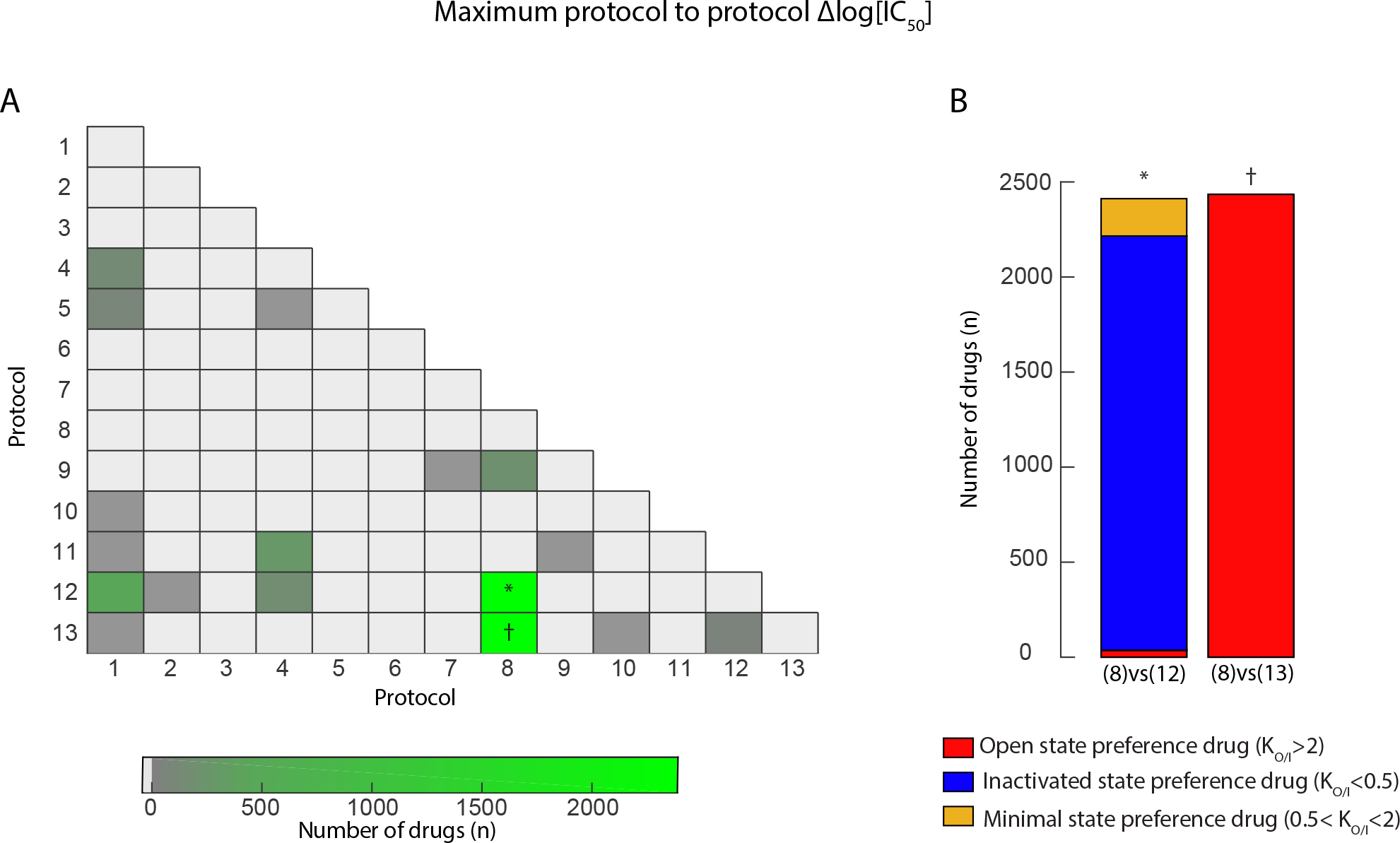
A) Heat map showing the frequency of protocol to protocol comparsion with the greatest difference in IC_50_ (max Δlog(IC_50_)) for each of the 6561 theoretical drugs. The highest frequency of max Δlog(IC_50_) was seen for comparison of protocols (8) vs (12) and (8) vs (13). B) Frequency of max Δlog(IC_50_) comparison of protocols (8) vs (12) and (8) vs (13) grouped by state binding preference. Inactivated state preference drugs and non-state preference drugs was most frequently observed in comparison (8) vs (12) (*) and max Δlog(IC_50_) for open state preference drugs was seen for comparison (8) vs (13) (†).

### Voltage protocol dependence of state occupancy

To better understand the mechanism behind these observations, we next examined how the state occupancy of the Kv_11.1_ channel varies with different voltage protocols. We examined the underlying Kv_11.1_ state occupancy during the 3 most prominent voltage protocols from Figure 4. Shown in Figure 5A are the voltage waveforms for the 0.1 Hz step-ramp protocol (protocol 8) and the 2 non-pulsed V_hold_ protocols at +20mV and −20mV (protocols 12 and 13 respectively). The respective time varying state occupancy for the open (red), closed (black) and inactivated (blue) states of the channel during each of these protocols is shown in Figure 5B. To compare the relative occupancy over time, for each protocol we calculated the state occupancy time-integral for each state expressed as a fraction of the total time integral, i.e., Fso_open_ for the open state, Fso_inact_ for the inactivated state and Fso_closed_ for the closed state (Figure 5C) (equation 6, *Materials and Methods*).

The relative occupancy of the open vs inactivated states (R_O/I_), for the different protocols is shown as a number above each protocol in Figure 5C. All the protocols we tested had a R_O/I_ < 1, indicating that there was a greater probability of channels spending time in the inactivated state than the open state. For the two non-pulsed protocols (Protocols 12 & 13) R_O/I_ decreased with increasing voltage (0.6 and 0.07 for −20 mV and +20 mV respectively), consistent with the known voltage dependent inactivation behavior of Kv_11.1_. The pulsed protocol (Protocol 8) had a close to equal time spent in the open and inactivated states with R_O/I_= 0.82. However, the pulsed protocol (protocol 8) dwelt predominantly in the closed state (Fso_closed_ 96.3%) in contrast to the non-pulsed protocols (Fso_closed_ 0.03% and 14.2% for protocols 12 & 13 respectively). In keeping with this, a lower occupancy of the open and inactivated vs closed states was evident for the pulsed protocol (Protocol 8) compared to non-pulsed protocols 12 & 13 (Figure 5C).

This data suggests that protocol 12, having the lowest R_O/I_, will be the most sensitive to inactivated state preference drugs, whereas protocol 8 will be almost equal in sensitivity to open state and inactivated state preference drugs. Moreover, the observation in Figure 4 that inactivated state preference drugs were most likely to have a maximum difference in affinity between protocols 8 & 12 can be attributed to the large difference in Fso_inact_. Similarly, the maximum difference in affinity for open preference drugs is seen between protocols 8 & 13 due to the large difference in Fso_open_ between these protocols.

**Figure 5.**
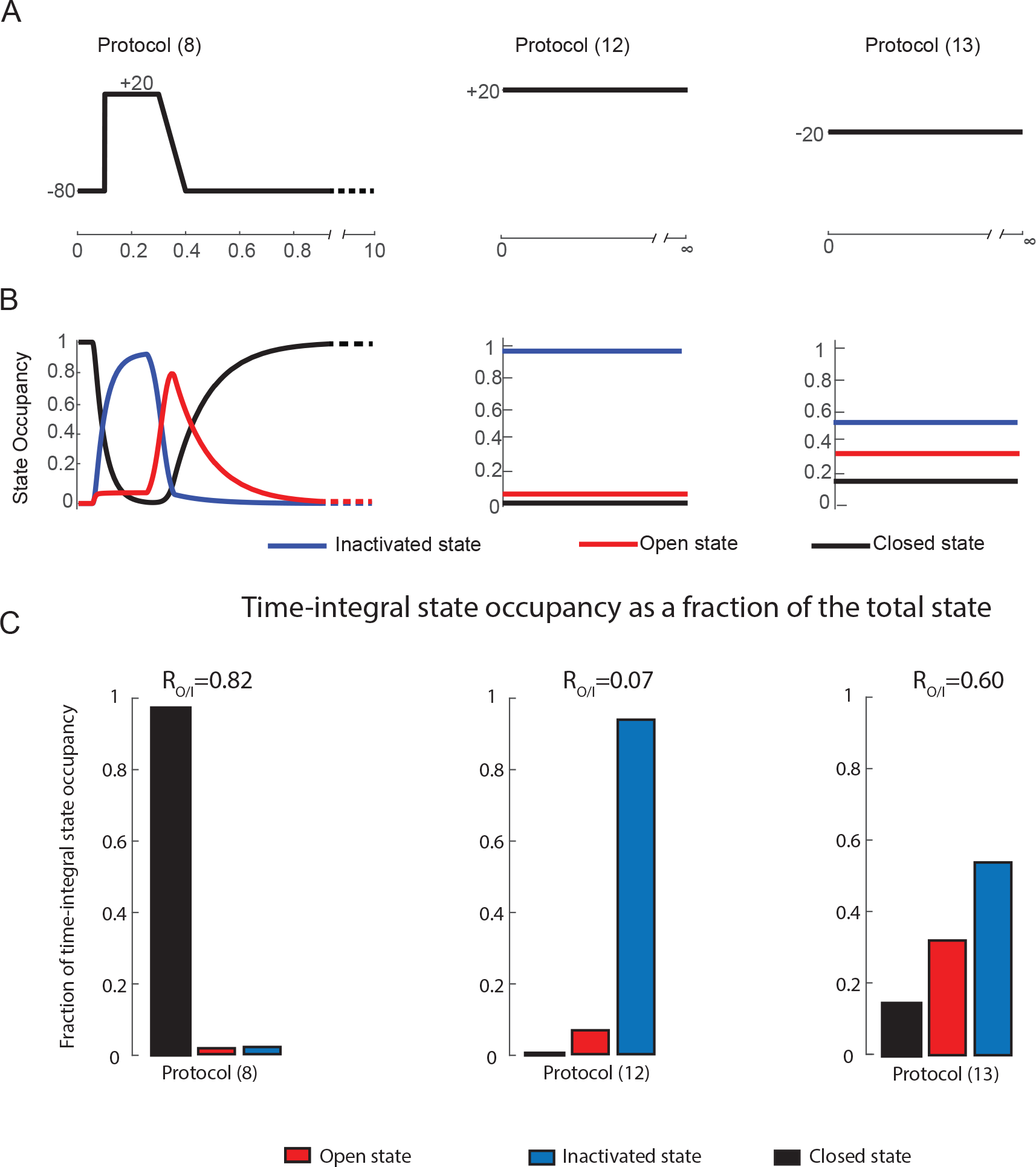
A) Voltage protocols for the subset of 3 of the 13 protocols with the greatest frequency of max Δlog(IC_50_) from Figure 4; used for *in silico* machine learning-training and validation. B) State occupancies for the open (red), inactivated (blue) and closed states (black) (shown as fraction of total state occupancy) for each of the 3 protocols depicted in panel A). Panels A & B are taken from Figure 1. C) Relative fraction of state occupancy (Fso) for the open (red) and inactivated (blue) and closed (black) states, expressed as a time-integral fraction or area-under-the-curve of the state occupancies shown in panel B. Relative state-occupancy (R_O/I_) of each protocol representing the ratio of Fso_open_ to Fso_inact_ is shown above each column graph.

### In silico prediction of state preferential drug binding characteristics

Our observations so far suggest that the state dependent binding properties of the drug and the gating state occupancy of the channel (as a function of the voltage protocol) interact in a predictable way to determine the differences in measured potency between protocols. To further test this hypothesis we sought to demonstrate the corollary of this; i.e. use measured differences in IC_50_ to predict a drug’s state preference. To do this we measured IC_50_ values for 2000 in silico drugs using protocols (8, 12 & 13) and from these created a set of 3 pairwise comparisons of ΔIC_50_. This set of pairwise comparisons was then used to train a neural network. A second set of 1000 different drugs was then used to test the accuracy of our predictions (Figure 6). Overall, the model performed extremely well, with a Pearson’s correlation coefficient between predicted and true state preference (K_O/I_) of R=0.930.

**Figure 6.**
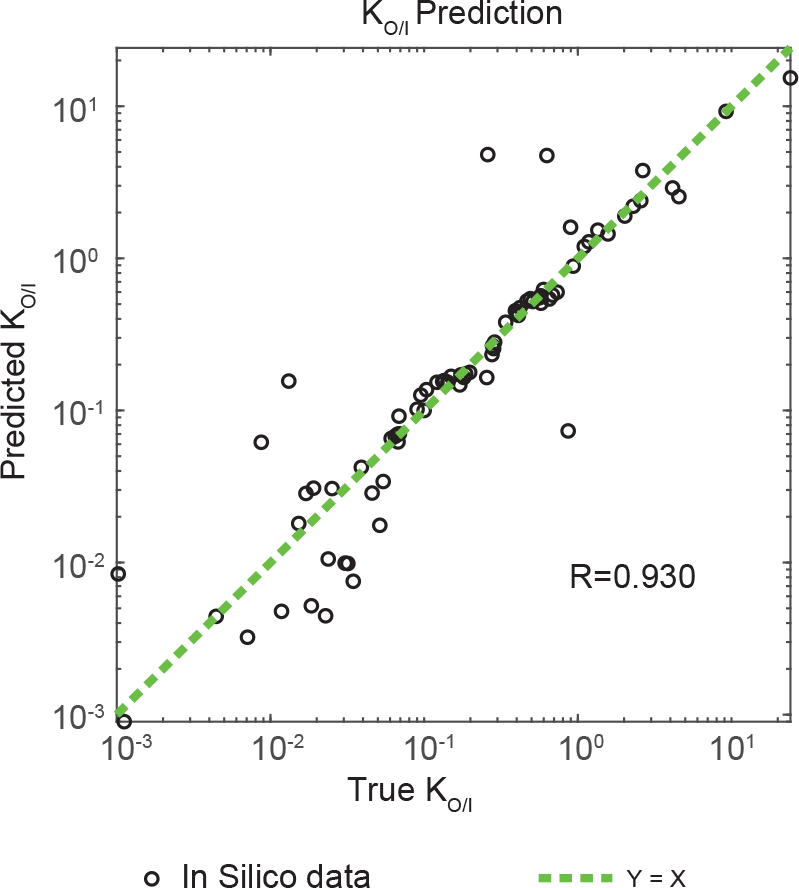
A *In silico* neural network algorithm was trained using IC_50_ values from Protocols (8, 12 and 13) to predict K_O/I_ for that drug. A separate *in silico* data set was used to independently validate the prediction algorithm (shown in black). An identity line (Y=X) is shown in dashed green. Accuracy of algorithm predictions is represented as the correlation coefficient R to the identity line.

We then sought to test whether the same 3 pairwise comparisons of ΔIC_50_ could be used to predict IC_50_ values for other voltage protocols. We were able to predict IC_50_ values in all cases (protocols 11, 1 and 10) with a high degree of accuracy (Figure 7 A-C). *In silico* predictions (shown in black) for protocols 1, 10 and 11 had a correlation coefficients of R= 0.984, R=0.988 and R=0.999 respectively.

**Figure 7.**
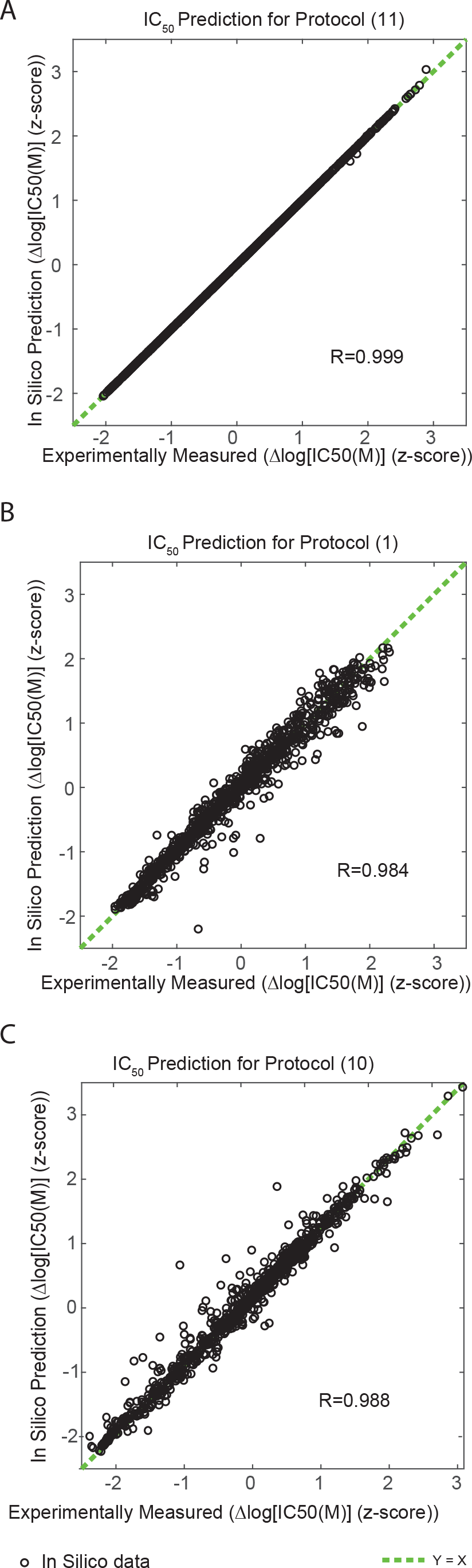
*In silico* neural network algorithms were trained using IC_50_ values from Protocols (8, 12 and 13) to predict IC_50_ values for other voltage protocols. An independent dataset using *in silico* data (black) was then used to validate the algorithm. Validation performance for each algorithm is shown in each of the three panels showing predictions for A) IC_50_ value for protocol (11), B) IC_50_ value for protocol (1) and C) IC_50_ value for protocol (10). An identity line (Y=X) is shown in dashed green. Accuracy of algorithm predictions is represented as the correlation coefficient R to the identity line for each prediction algorithm. The predicted and experimentally measured IC_50_ values are shown as the z-score transformed Δlog[IC_50_].

### In vitro measurement of protocol dependence of IC50

We next tested whether drugs with varying state preference also showed different measured potencies in manual patch clamp electrophysiology assays, depending on the voltage protocol used. Typical Kv11.1 current traces recorded using six of the protocols examined *in silico* (1, 8, 10, 11, 12, 13) in response to increasing [cisapride] are shown in Figure 8. To highlight the differences in measured potency between protocols, the 60nM dose of cisapride is highlighted red for each case. For the short-pulse protocols (8 & 10), 60 nM cisapride blocked measured current by ~25 %. In comparison, the same concentration caused 75-80 % reduction in current amplitude when measured using the long-pulse protocol (Protocol 1) or the non-pulse protocols (11, 12 & 13).

**Figure 8.**
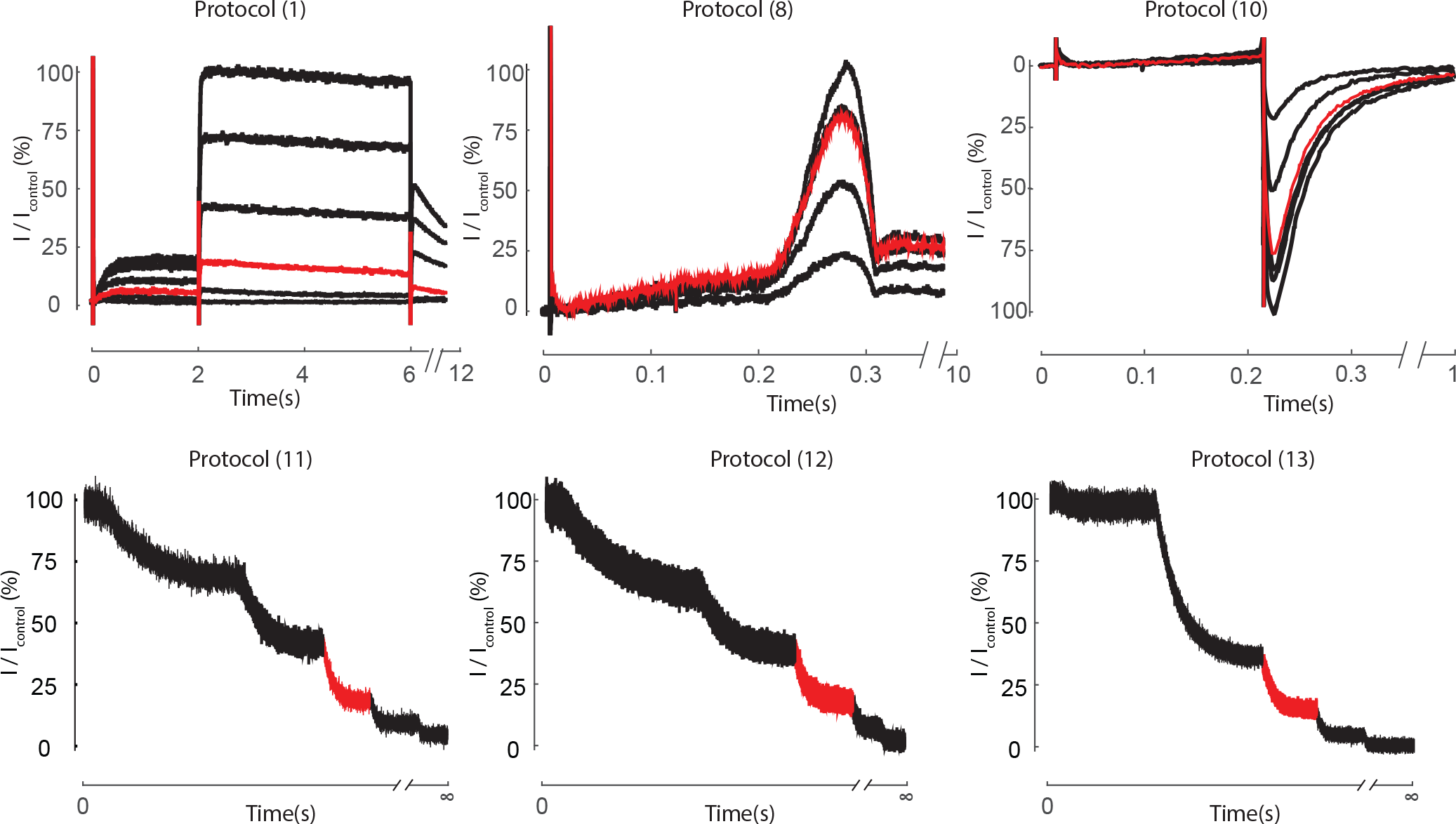
*In vitro* experimental data of evoked I_Kr_ currents for six voltage protocols using CHO cells stably expressing Kv11.1. Kv11.1 currents were measured whilst cells were perfused with bath solution in 0, 10, 20, 60, 200 and 600nM cisapride. Highlighted in red is the I_Kr_ current perfused with 60nM cisapride for each of the six protocols demonstrating protocol to protocol variation in the degree of Kv11.1 block with the same dose of drug concentration.

The same set of *in vitro* experiments were also carried out for three other drugs (clozapine, verapamil and terfenadine), dose response curves constructed for each of the six protocols (Figure 6A-D – left side panels), and IC_50_ values derived from fits of the Hill equation (Figure 6A-D – right side panels). A maximum of 2.79, 8.32, 1.93 & 3.09 – fold

ΔIC_50_ was measured between protocols for verapamil, cisapride, clozapine and terfenadine respectively. Overall, significant differences between the IC_50_ measured using the six protocols were observed for verapamil, cisapride and terfenadine (P = 0.0297, 0.0005 and 0.001 respectively; ANOVA), but not for clozapine (P= 0.3406). This result suggests that verapamil, cisapride and terfenadine may have state preferential binding properties. Furthermore, verapamil and cisapride demonstrated a higher affinity when measured with protocol 12 compared to protocol 8 (1.77-fold and 5.96-fold ΔIC_50_, respectively), consistent with our *in silico* model observations for inactivated state preference drugs (Figures 2 and 3A).

**Figure 9.**
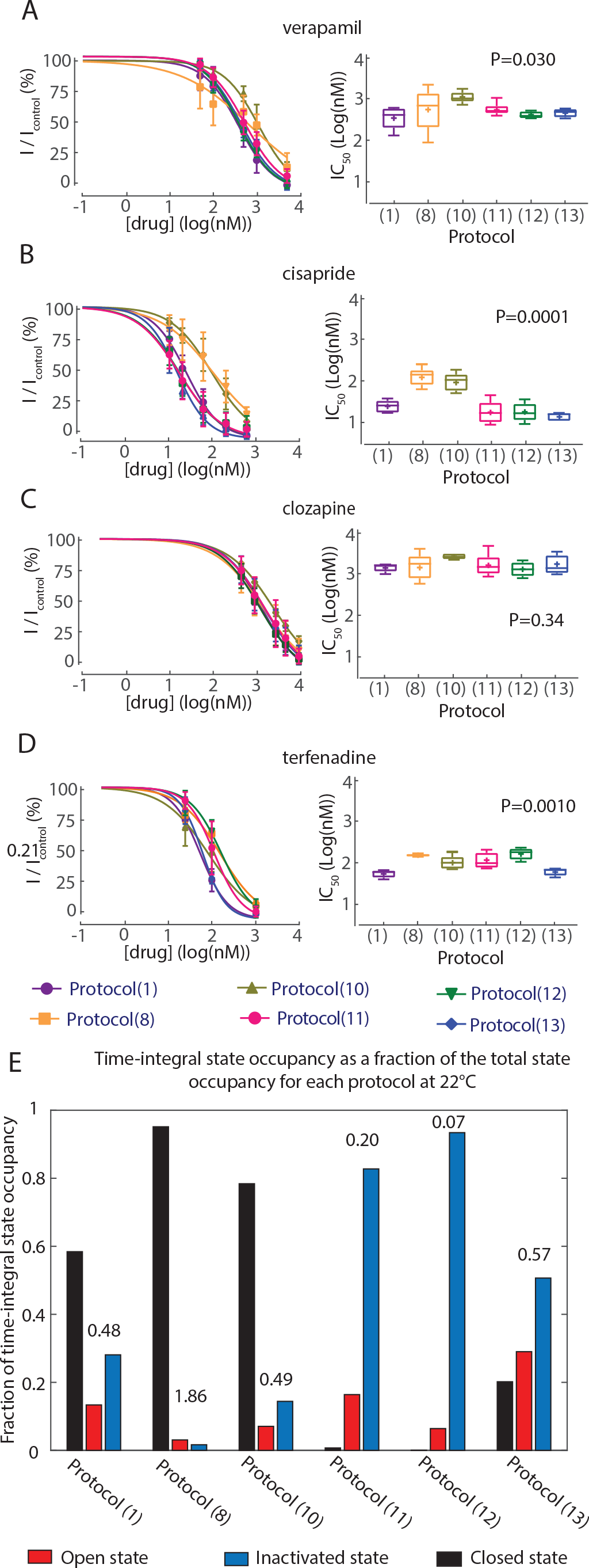
*In vitro* experimental data. A) Examples of *in vitro* Kv_11.1_ currents evoked when voltage clamped using voltage protocols 8 (left) and 12 (right) in the presence of increasing concentrations of drug. A – D) Dose response curves (left) and box-plot of corresponding IC_50_ values for each of the 6 voltage protocols (1, 8, 10, 11, 12, 13) (n=4 – 6) in the presence of verapamil, cisapride, clozapine and terfenadine respectively. The central line shows the median value of each protocol. The mean for each box is depicted as a (+). Box-plot boxes extend to the 25^th^ and 75^th^ percentile of the data for each protocol and the whiskers show the 95% confidence intervals for IC_50_ values for each protocol. P values for analysis of variance tests (ANOVA) are shown for each drug. E) Relative proportions of Fso for each of the 6 voltage protocols, demonstrating underlying differences in state occupancy at 22 °C. Relative state-occupancy (R_O/I_) of each protocol representing the ratio of Fso_open_ to Fso_inact_ is shown above each column in the figure.

To understand these observed differences in IC_50_ we again considered the channel state occupancy during the individual voltage protocols. For consistency between *in vitro* experiments and *in silico* state occupancy calculations, the state occupancies in Figure 9E were calculated for 22 °C by scaling the transition rates in the Markov model (Figure 1) for temperature (See Methods). For cisapride, there is a correlation between measured potency and Fso_inact_, i.e. lower IC_50_s were measured for protocols with greater Fso_inact_ (Protocols 11, 12 and 13), and highest IC_50_s were measured for protocols with lowest Fso_inact_ (Protocol 8), consistent with preferential binding to the inactivated state. In addition to this, amongst the pulsed protocols, block by cisapride is less potent when measured with the short-pulsed protocols (8 & 10); which have lower Fso_open_ and Fso_inact_ compared to the long-pulsed protocol (1) (Figure 9E). This suggests that there is a requirement for a longer channel opening time in order for the drug bound states (either open or inactive) to reach their true equilibrium. Overall, a similar pattern was observed for verapamil, though to a lesser degree (Figure 9A).

For terfenadine, while there are significant differences in potency of block measured between protocols, the correlation between IC_50_ and Fso_inact_ is not apparent. In particular the relationship is disrupted by the downward shift in IC50 measured using protocols 8 & 10 (as compared to cisapride for example). We note that these protocols have a greater proportion of Fso_closed_, suggesting that drug remaining bound to the closed state, or drug trapping may be a modulating factor in the measured in IC_50_ for terfenadine. Finally, for clozapine, the potency of block was the same regardless of the state occupancy of the voltage protocol used (Figure 9C).

### In silico prediction of in vitro drug binding characteristics

Since there appears to be a qualitative agreement between state occupancy and measured potency for most of the drugs tested, we next sought to use our *in silico* trained prediction algorithm (Figure 6) to predict the state preference (K_O/I_) of the 4 drugs using the *in vitro* measurements of IC_50_ from protocols 8, 12 & 13. K_O/I_ predictions for verapamil, cisapride, clozapine and terfenadine are shown in Table 1.

**Table 1:**
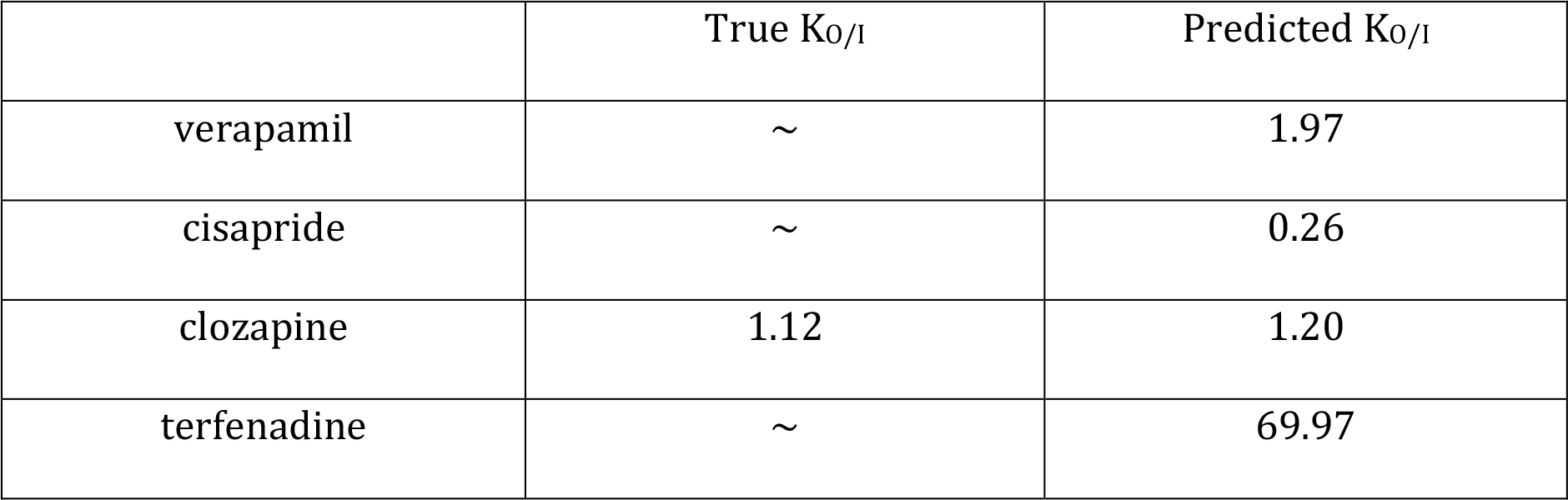
True K_O/I_ compared to predicted K_O/I_ for the 4 *in vitro* drug examples. Note only clozapine has a fully parameterized model of binding to Kv_11.1_ from Hill et al. 2014, and therefore the only drug that has an *in vitro* validated value for K_O/I_.

| | True $K_{0/I}$ | Predicted $K_{0/I}$ |
| --- | --- | --- |
| verapamil | ~ | 1.97 |
| cisapride | ~ | 0.26 |
| clozapine | 1.12 | 1.20 |
| terfenadine | ~ | 69.97 |

Similarly, we used the algorithms from Figure 7 to predict the measured IC_50_ values of other *in vitro* voltage protocols (1, 10 & 11). Predictions for our four *in vitro* drugs (verapamil, cisapride, clozapine and terfenadine) are shown in Figure 10 in red. To put these *in vitro* predictions in context, we added the predictions of our 1000 theoretical *in silico* drugs from Figure 7 in grey. A similar prediction accuracy was achieved for the four *in vitro* drug predictions (shown in red) as for the *in silico* drug predictions. Notably, *in vitro* IC_50_ prediction algorithms were better for the non-pulse protocol (Protocol 11, Figure 10A) and the long-pulse protocol (Protocol 1, Figure 10B), compared to the short-pulse protocol (Protocol 10, Figure 10C). Overall, our accuracy of *in vitro* IC_50_ prediction using our *in silico* prediction algorithm, was within one logarithmic unit of the measured *in vitro* IC_50_ in all but one case. Details of *in vitro* measured and *in silico* predicted IC_50_‘s, respective Hill coefficients as well as K_O/I_ are shown in Supplementary Table 1.

**Figure 10.**
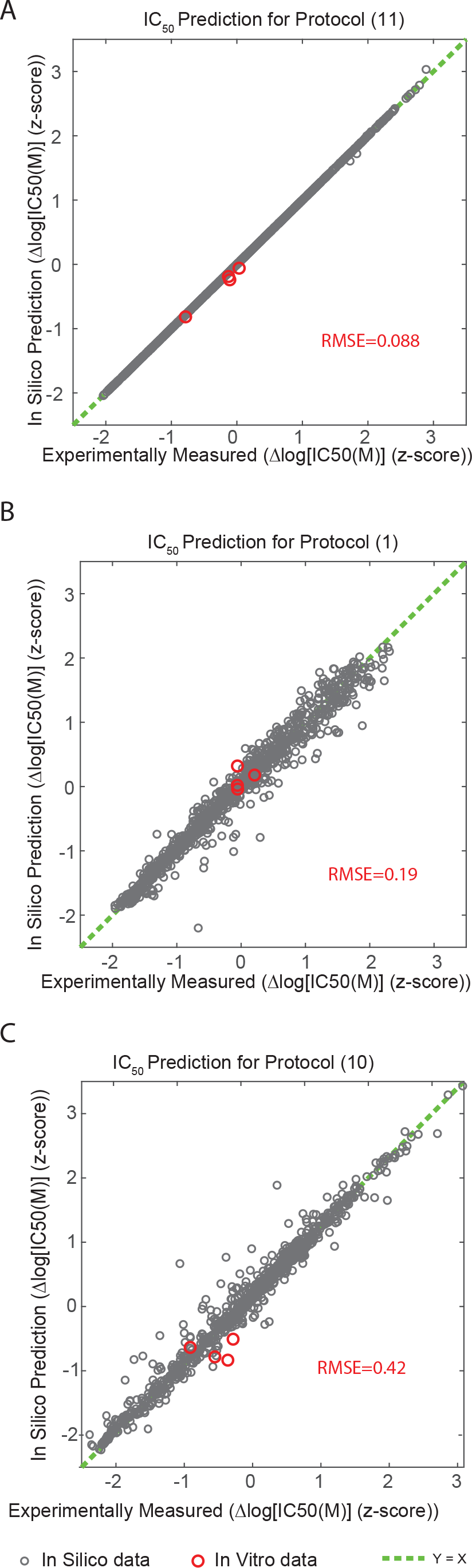
*In silico* neural network algorithms were trained using *in silico* IC_50_ values from protocols (8, 12 & 13) to predict *in vitro* IC_50_ values for other voltage protocols using *in vitro* measurements of IC_50._. For each panel, predictions vs true measured IC_50_s for the 4 *in vitro* drugs (verapamil, cisapride, clozapine and terfenadine (shown in red). For context, *in silico* predictions from Figure 7 are shown in grey. A) IC_50_ value for protocol (1), B) IC_50_ value for protocol (10) and C) IC_50_ value for protocol (11). An identity line (Y=X) is shown in dashed green. The accuracy of *in vitro* prediction for each of the three voltage protocols is represented as the root mean squared error (RMSE) for each panel.

## Discussion

In this study, we have shown that state-dependent drug binding to Kv_11.1_ channels is a significant factor in determining the range of IC_50_s measured for individual compounds using different voltage protocols. Furthermore, these differences occur in a predictable manner as a function of the state occupancy of the channel. As a result, both a drug’s state preference, as well as its potency measured using other voltage protocols can be inferred from simple equilibrium measures of block.

### Factors contributing to protocol-dependent potency of block

The variability in potency of Kv_11.1_ block measured using different voltage protocols has been reported in several studies (Kirsch *et al.*, 2004; Yao *et al.*, 2005; Milnes *et al.*, 2010). For example, the IC_50_ for cisapride varies over a 60 fold range, depending on the protocol used (Potet *et al.*, 2001; Rezazadeh *et al.*, 2004). Our data demonstrate that the synergy of the state preference in drug binding as well as voltage protocol-dependent state occupancy results in such observed variations in affinity. Therefore, drugs that preferentially bind to a specific gating state will have a higher measured affinity for Kv_11.1_ when using a voltage protocol that favors that gating state, and vice versa. In contrast, for drugs that have no state preference there is no dependence of the measured IC_50_ on the voltage protocol used. However, the data in Figure 3 demonstrates that variations in IC_50_ can also occur regardless of state preference, suggesting that the kinetics of drug binding can also have an impact on measured potency. Drugs that block Kv_11.1_ require channel opening to gain access to their binding site in the inner cavity of the channel pore (Ficker *et al.*, 1998; Walker *et al.*, 1999). Therefore drugs with slow binding kinetics will have a relatively lower affinity when measured using protocols where occupancy of the open/inactivated states is short (relative to the on rate for drug binding), and have a relatively higher apparent affinity when measured using voltage protocols with long depolarization steps or non-pulsed protocols that promote prolonged occupancy of the open and inactivated states. In our *in vitro* data, this phenomenon is particularly evident for cisapride (Figure 9B) which has a lower potency when measured using short-pulse protocols (protocols 8 & 10) where channel open times are on the order of hundreds of milliseconds. This is consistent with previous publications measuring the timecourse of block of Kv11.1 by cisapride with time constants of ~ 20 s (Windley *et al.*, 2016; 2017).

Another factor which may modify a drugs measured potency in a protocol dependent manner is trapping, whereby upon closing of the channel in response to membrane repolarisation, a drug is unable to dissociate from the channel cavity (Mitcheson *et al.*, 2000). As a result, accumulation of block can occur for trapped drugs over successive sweeps of a protocol with a high proportion of Fso_closed_ (Li *et al.*, 2017; Windley *et al.*, 2017). Conversely, non-trapped drugs dissociate during the closed intervals resulting in a lower measured potency. The differences in our *in vitro* IC_50_s for terfenadine and cisapride are consistent with this. The potency of block for cisapride, a non-trapped drug (Li *et al.*, 2017; Windley *et al.*, 2017), measured using protocols 8 & 10 (with Fso_closed_ of 95 % and 78 % respectively) is less than when measured using all other protocols which have with lower Fso_closed._ Conversely, for terfenadine, a trapped drug (Kamiya *et al.*, 2008; Windley *et al.*, 2017), the potency measured with protocols 8 & 10 is increased relative to the other protocols – consistent with accumulation of block in the closed state. This ability to identify truly trapped drugs, as opposed to those such as cisapride that display ‘virtual trapping’ as a result of slow unbinding kinetics (Windley *et al.*, 2017), is important since it has been suggested that ‘true’ trapping may confer an additional arrhythmia risk (Di Veroli, Davies, H Zhang, Abi-Gerges, and Boyett, 2013a).

We also noted some variation in the Hill coefficients measured across the various *in vitro* protocols (see Supplemental Table 1). While most of the coefficients were close to 1, consistent with previously published reports, there were some exceptions. For example, a lower Hill coefficient was found for verapamil using protocol 8 (Hill coefficient = 0.55), while in general Hill coefficients for terfenadine were slightly higher than 1 (eg. The highest Hill coefficient for terfenadine was 1.67 for protocol 13). These observations can perhaps be explained by factors such as the difficulty in accurately measuring the degree of block for some drugs with certain protocols. For example, the kinetics of binding of terfenadine to Kv11.1 have previously been shown to be problematic for measuring steady state block, since the timecourse of block is so slow (Windley *et al.* 2016). In our experiments, an underestimation of the degree of block at low concentrations, when the timecourse of terfenadine block is slowest, would explain the Hill coefficients being greater than 1.

### Predicting potency across different voltage protocols

Defining the potency of Kv_11.1_ block is an important part of existing preclinical safety screening, where a safety index value is determined by comparing the inhibitory concentration of Kv_11.1_ (IC_50_) to the maximum plasma concentration (C_max_) of the drug (Redfern *et al.*, 2003). However, the degree of protocol-dependence of measured potency that we and others have demonstrated (Supplement Table 2-5), makes this is problematic because: 1) A single affinity measure does not represent a ‘true’ affinity for Kv_11.1_; it only represents a drug’s affinity for one specific voltage protocol; and 2) Even if a single voltage protocol could be agreed upon to standardize drug affinity assays, it would not adequately reflect the wide ranging effects of external conditions such as heart rate, QT prolongation, and hypokalaemia. Our data shows that there is a predictable relationship between IC_50_s measured using different protocols. Furthermore, using IC_50_s measured from just a few simple voltage protocols, it is possible to predict what a drugs potency will be when measured using other voltage protocols. For *in silico* data, this can be done with a very high degree of accuracy (Figure 7), whilst for *in vitro* data, the degree of accuracy is reduced, perhaps as a result of factors such as drug trapping, which are not implicit in the Markov model used in this study. Nevertheless, this demonstration of a quantifiable relationship between the variable measured potencies of Kv_11.1_ block is important as it provides a framework for direct comparison of potencies measured between groups using different assays, and may allow for reinterpretation of legacy datasets obtained using different protocols.

### Prediction of state-dependent binding characteristics

It has previously been shown that high affinity binding to Kv_11.1_ occurs, at least for some drugs, because of preferential binding to the inactivated state (Suessbrich *et al.*, 1997; Ficker *et al.*, 1998; Numaguchi *et al.*, 2000; Perrin, Kuchel, *et al.*, 2008). Furthermore, preferential binding to the inactivated state has been associated with increased prolongation of the action potential duration (Lee *et al.*, 2016), and several drugs that have been withdrawn from market (eg. astemizole and terfenadine) or had their use severely restricted (eg. cisapride), preferentially bind to the inactivated state (Perrin, Kuchel, *et al.*, 2008). However, quantification of state-dependent binding is difficult, since it cannot be readily measured experimentally. Rather, indirect measures relying on changes in affinity that occur as a result of perturbation of the inactivated state using site directed mutagenesis (Suessbrich *et al.*, 1997; Ficker *et al.*, 1998; Numaguchi *et al.*, 2000; Perrin, Kuchel, *et al.*, 2008), changing potassium concentration (Kamiya *et al.*, 2008), or measurement of drug binding kinetics (Hill *et al.* 2014) are routinely used.

In this study, we show that a machine learning algorithm, trained using IC_50_s measured from just three simple voltage protocols, can accurately determine the degree of state preference drugs *in silico* (Figure 6). We also used the same algorithm to predict the state preference of four drugs – clozapine, cisapride, verapamil and terfenadine - based on our *in vitro* measured IC_50_s. For clozapine, the only drug we tested for which there is a published, fully parameterized *in silico* model (Hill *et al.*, 2014), meaning the state dependent binding properties are unambiguously known, the algorithm correctly identified this as a drug with no state preference (Table 1). For cisapride, the algorithm predicted inactivation state preference (K_O/I_ = 0.26), consistent with previously published studies using mutagenesis, which showed a ratio of 0.37 between the IC_50_ for wildtype Kv_11.1_ and the N588K inactivation deficient mutant Kv_11.1_ (Perrin, Kuchel, *et al.*, 2008). The data in the literature for verapamil is mixed. Previously publications show a low variation in measured IC_50_ values from any 2 protocols (maximum 6-fold difference; Supplement Table 2), suggesting verapamil has weak or non-preferential state binding to Kv_11.1_. This is consistent with our model prediction of K_O/I_ =1.97. However, other studies have shown that potency of block of the S620T inactivation deficient Kv_11.1_ mutant by verapamil was less potent than for wildtype (S Zhang *et al.*, 1999) – consistent with inactivation state preference. However, this mutation has also been shown to have a gating-independent effect on drug binding (Guo *et al.*, 2006), i.e. mutation at this residue directly disrupts the drug binding site, which might confound the interpretation of the effects of this mutant in relation to state preference. Finally, our predicted K_O/I_ for terfenadine was 69.67 suggesting a strong preference for binding to the open state of Kv_11.1_. This is inconsistent with published data using either mutagenesis (Perrin, Kuchel, *et al.*, 2008) or potassium inhibition of inactivation (Kamiya *et al.*, 2008) that identified terfenadine as a drug that preferentially binds to the inactivated state. One explanation for this is that terfenadine exhibits closed state trapping (Kamiya *et al.*, 2008; Windley *et al.*, 2017), which is not accounted for in the Markov model we used, that confounds the *in vitro* predictions for this drug.

## Limitations

The main limitation of this study, in relation to prediction of *in vitro* data using an algorithm developed using *in silico* data is the characteristics of the available Markov models describing drug binding to Kv11.1 - particularly in relation to drugs that exhibit closed state trapping. While there are models in the literature that include either state-dependent binding (Di Veroli, Davies, H Zhang, Abi-Gerges, and Boyett, 2013b; Lee *et al.*, 2016) or trapping (Di Veroli, Davies, H Zhang, Abi-Gerges, and Boyett, 2013a; Li *et al.*, 2017), there are no models that include both. Development of a more complete Markov description of drug binding which included both state-dependent binding and trapping would further improve predictions based on *in vitro* data and indeed, the quantification of state dependent binding enabled using the approach reported here will likely facilitate the parameterization of such models.

Secondly, it is well known that the kinetics of drug binding to Kv11.1 channels (Windley et al. 2016) as well as the kinetics of Kv11.1 channel gating (Vandenberg et al. 2006) are temperature dependent. This is evident in the differences that we observe in the I_Kr_ current at 37°C *in silico* (Figure 1B) and 22°C *in vitro* (Figure 8). Ideally therefore, an *in vitro* dataset gathered at 37°C would be desirable for further validation. Despite this, we still achieve good prediction accuracy with our machine learning models both for state-preference and drug affinity. This is consistent with previous published data showing that even though the kinetics of drug binding to Kv11.1 for the 4 drugs examined here are temperature dependent, the IC_50_s for block are not (Windley et al. 2016, Windley et al. 2018, Hill et al. 2014).

## Conclusion

In this study, we have demonstrated that state-dependent drug binding is a major factor in determining the potency of Kv_11.1_ block measured using different protocols, with the exact measured potency also affected by factors such as drug binding/unbinding kinetics and closed state trapping. As a result, we show that inter-protocol differences in measured IC_50_ occur in a predictable way, meaning a compound’s potency measured using any voltage protocol can be reliably estimated from knowledge of the channel state occupancy. This is an important step since it allows for direct comparison of potencies measured using different assays, as well reinterpretation of legacy datasets obtained using different protocols as part of previous compound screens. Furthermore, we also show that a drug’s state preference can be inferred from differences in its IC_50_ measured from commonly used protocols that are simple enough to be readily deployed on high throughout, automated patch clamp platforms. This is the first demonstration of quantification of state preference without the need for either measuring kinetics of block/unblock, or disruption of Kv11.1 inactivation (using mutagenesis or ionic concentration). The ability to simply quantify state-preference will facilitate the development of more complete models of drug binding to Kv_11.1_ and improve our understanding of proarrhythmic risk associated with Kv_11.1_ blocking compounds.

## Acknowledgements

The authors thank, Mark Hunter, Dr Stewart Heitmann, and Dr Melissa Mangala for informative discussion.

## Authorship Contributions

Participated in research design: Lee, Vandenberg, Hill Conducted experiments: Lee, Windley, Perry

Contributed to new reagents or analytical tools: Lee, Hill, Vandenberg Performed data analysis: Lee, Hill

Wrote or contributed to the writing of the manuscript: Lee, Windley, Perry, Vandenberg, Hill

## Footnotes

This work was supported by grants from the National Health and Medical Research Council of Australia [APP1088214]. JIV is supported by an NHMRC Senior Research Fellowship [APP1019693]. WL is supported by a National Heart Foundation of Australia Health Professional Scholarship [APP101552].

## Abbreviations

hERG: human ether-a-go-go related gene
*I*_Kr_: rapid component of delayed rectifier current
aLQTS: acquired Long QT syndrome
TdP: Torsade de Pointes
C_max_: maximum therapeutic plasma concentration
CiPA: Comprehensive *In vitro* Proarrhythmic Assay
APD: action potential duration
ΔIC_50_: difference in log [IC_50_] measured between two protocols
Fso: Fractional state occupancy
R_O/I_: Ratio of the Fractional state occupancy of the open state vs the inactivated state

