## Supplemental data for "Protocol-dependent differences in IC_50_ values measured in hERG assays occur in a predictable way and can be used to quantify state preference of drug binding"

FIGURE 1

*A.*

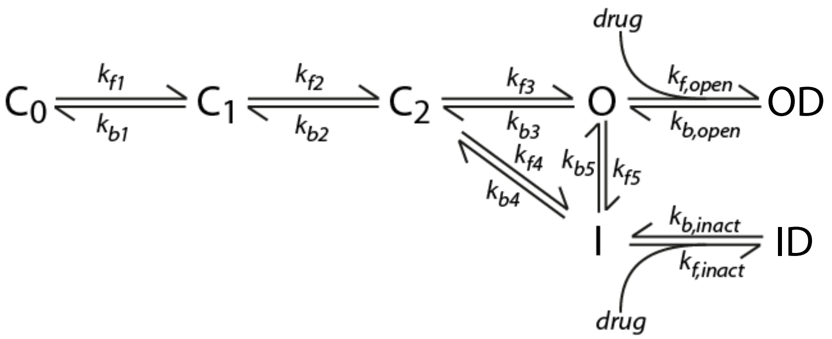


*B.*

| | $\alpha_0$ | $z_\alpha$ | $\beta_0$ | $z_\beta$ |
|---|---|---|---|---|
| 1: $C_0 - C_1$ | 0.1161 | 0.299 | 0.2442 | -1.604 |
| 2: $C_1 - C_2$ | 0.1235 | 0 | 0.1911 | 0 |
| 3: $C_2$ – O | 0.0578 | 0.971 | 0.000349 | -1.062 |
| 4: $C_2$ – I | 0.000052 | 1.525 | 0.0000000085 | -1.842 |
| 5: O – I | 0.2533 | 0.5953 | 0.0522 | -0.8209 |

**Supplementary Figure 1.** A) Markov state model of Kv11.1-drug interaction. Parameters describing voltage-dependent Kv11.1 gating transitions are presented in the table in (B). Drug binding and unbinding is described as forward and backward rate constants kf,open and kb, open respectively for the open state and kf, inact and kb, inact for the inactivated state. These drug binding and unbinding rate constants are specified for each individual drug scenario.

Protocol (1)

Voltage Waveform

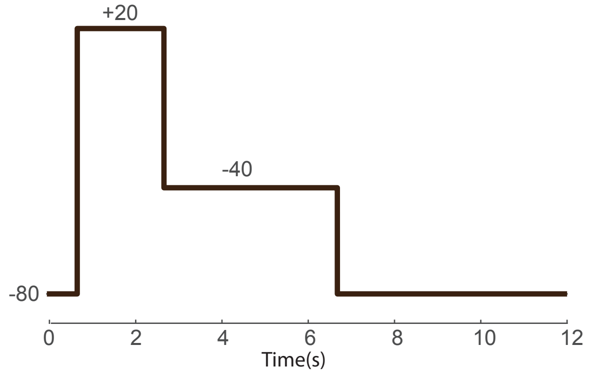


In Silico Kv11.1 Current

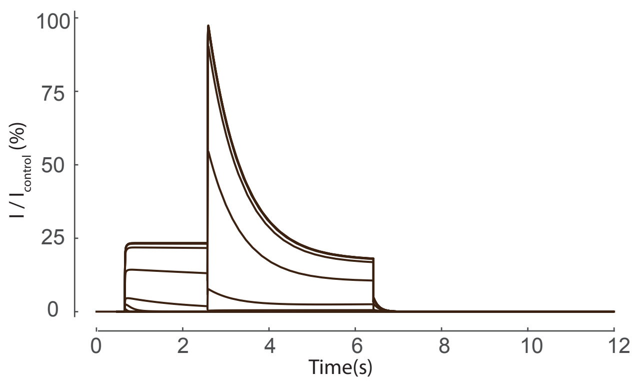


In Vitro Kv11.1 Current

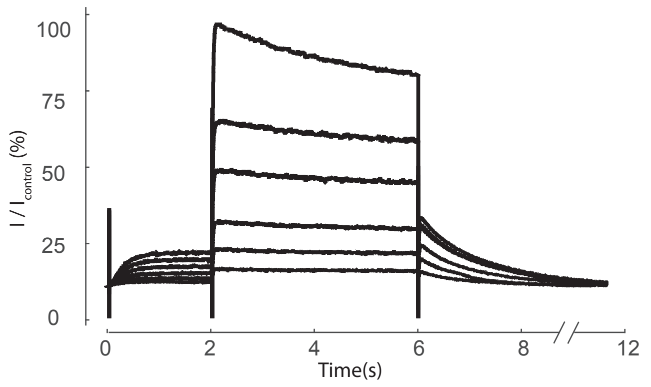

### Protocol (2)

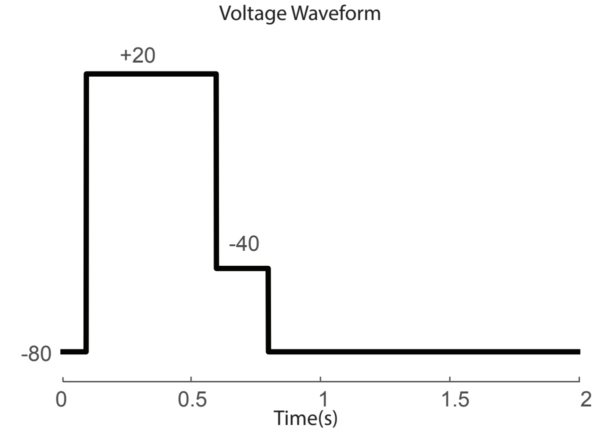
Voltage Waveform
+20
-40
-80
0
0.5
1
1.5
2
Time(s)

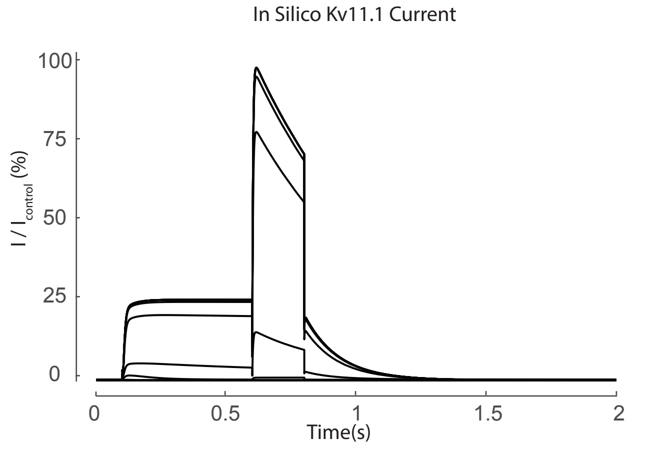
In Silico Kv11.1 Current
100
75
50
25
0
$I / I_{control}$ (%)
0
0.5
1
1.5
2
Time(s)

### Protocol (3)

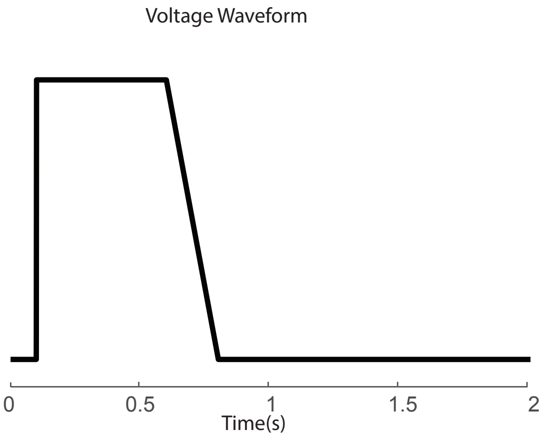
Voltage Waveform
0
0.5
1
1.5
2
Time(s)

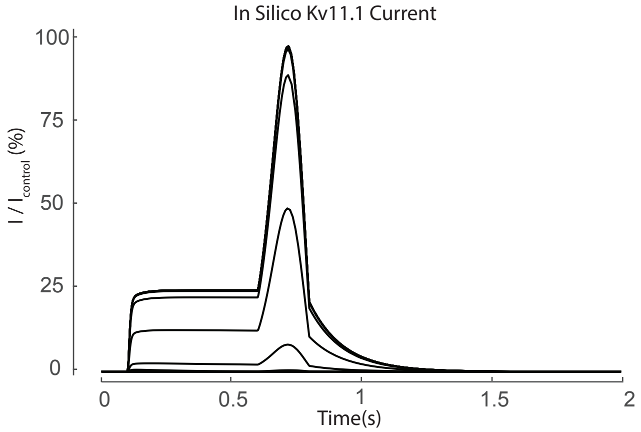
In Silico Kv11.1 Current
100
75
50
25
0
I / I_control (%)
0
0.5
1
1.5
2
Time(s)

Protocol (4)

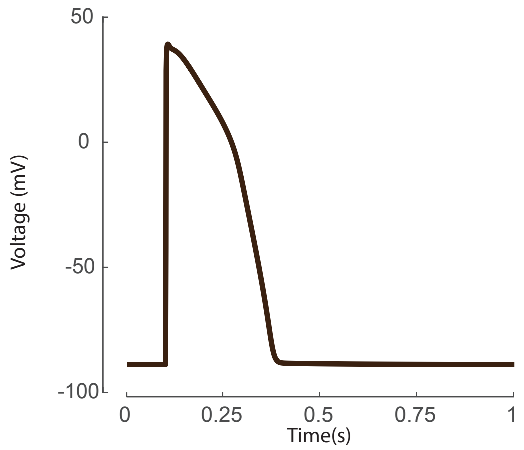
50
0
-50
-100
Voltage (mV)
0
0.25
0.5
0.75
1
Time(s)

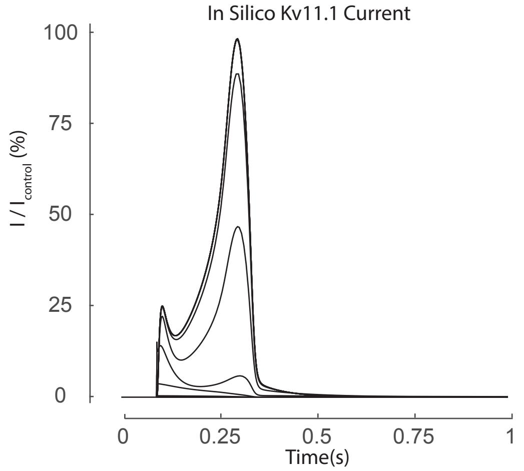
In Silico Kv11.1 Current
100
75
50
25
0
I / I$_{control}$ (%)
0
0.25
0.5
0.75
1
Time(s)

### Protocol (5)

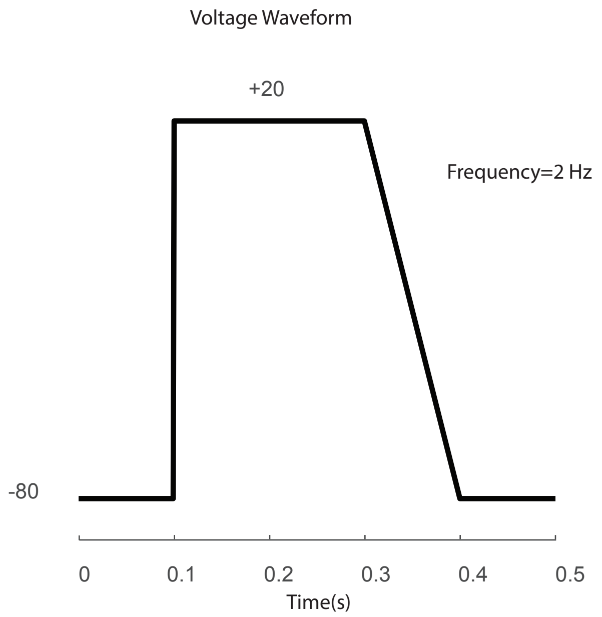
Voltage Waveform
+20
Frequency=2 Hz
-80
0
0.1
0.2
0.3
0.4
0.5
Time(s)

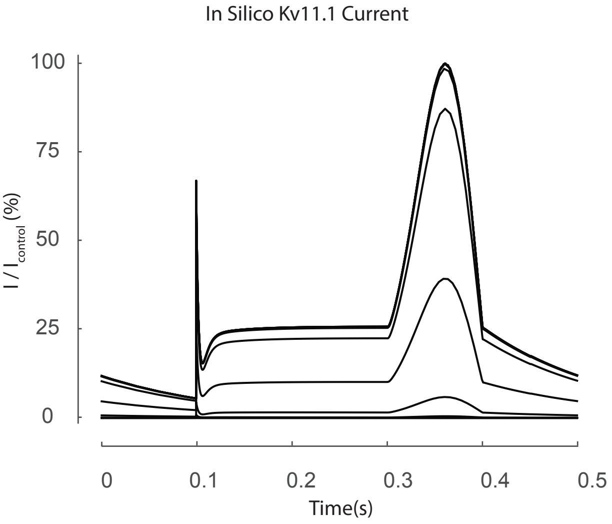
In Silico Kv11.1 Current
100
75
50
25
0
I / I control (%)
0
0.1
0.2
0.3
0.4
0.5
Time(s)

### Protocol (6)

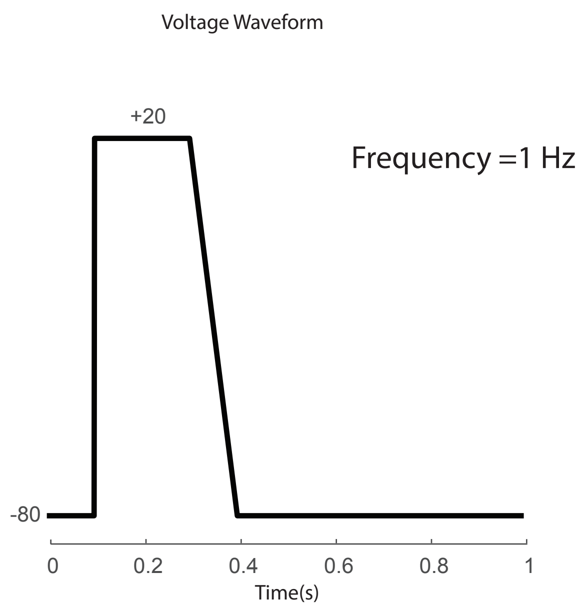
Voltage Waveform
+20
Frequency =1 Hz
-80
0
0.2
0.4
0.6
0.8
1
Time(s)

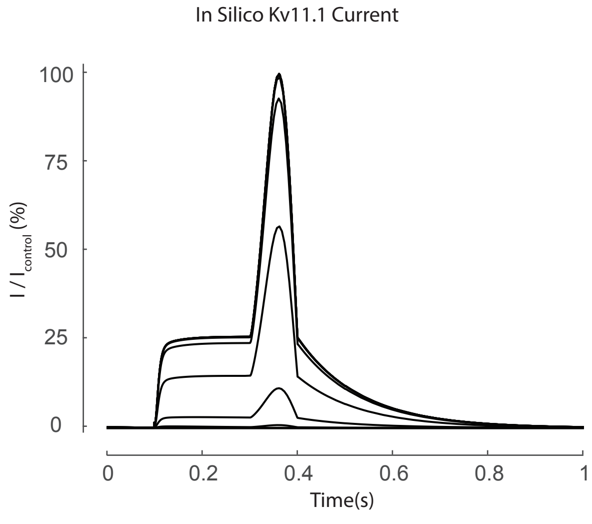
In Silico Kv11.1 Current
100
75
50
25
0
I / I control (%)
0
0.2
0.4
0.6
0.8
1
Time(s)

### Protocol (7)

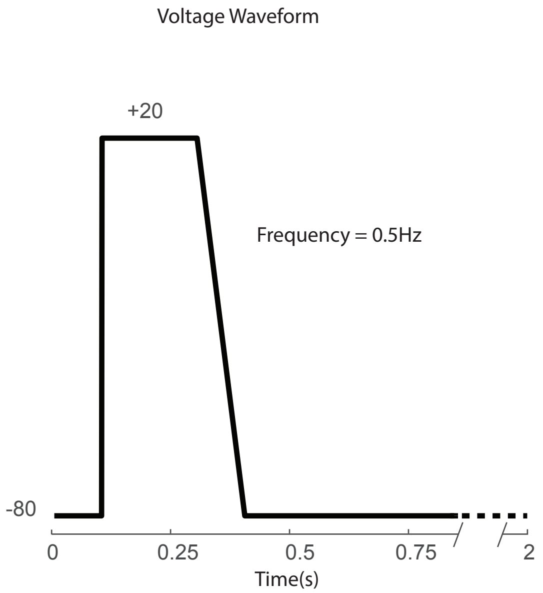
Voltage Waveform
+20
Frequency = 0.5Hz
-80
0
0.25
0.5
0.75
2
Time(s)

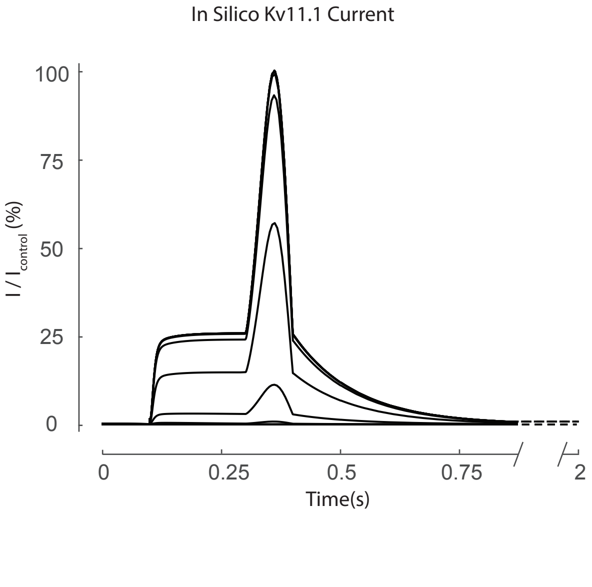
In Silico Kv11.1 Current
100
75
50
25
0
I / I control (%)
0
0.25
0.5
0.75
2
Time(s)

### Protocol (8)

Voltage Waveform

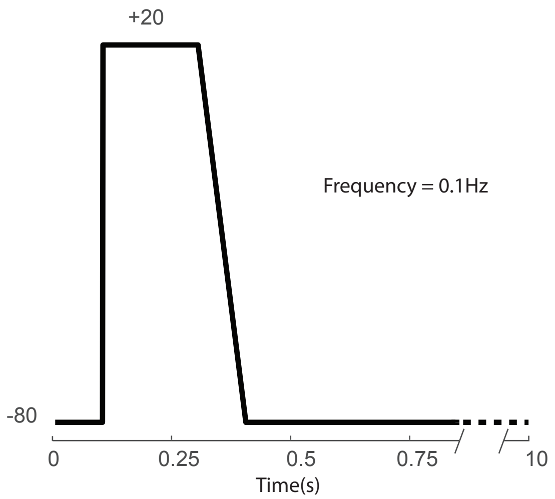


In Silico Kv11.1 Current

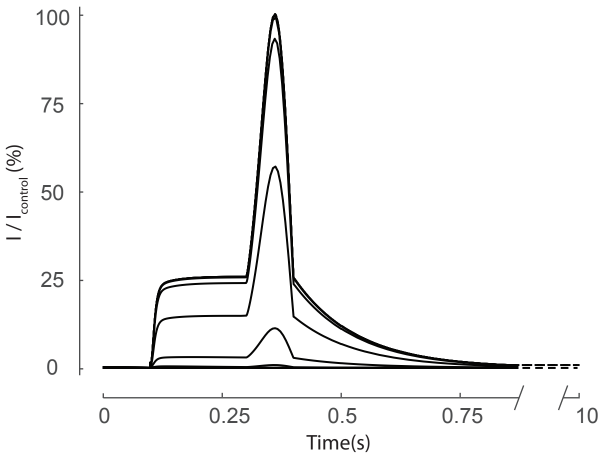


In Vitro Kv11.1 Current

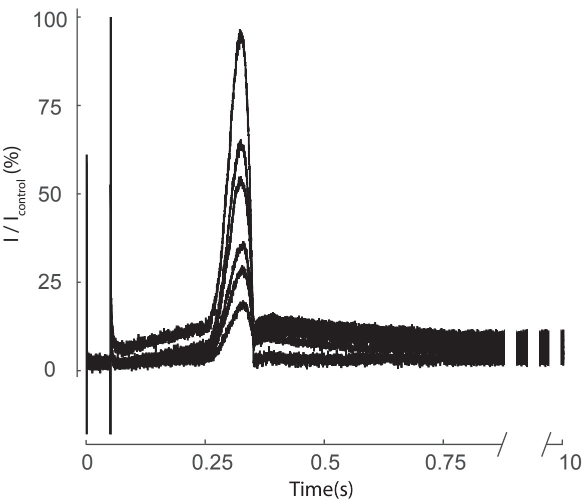

### Protocol (9)

Voltage Waveform

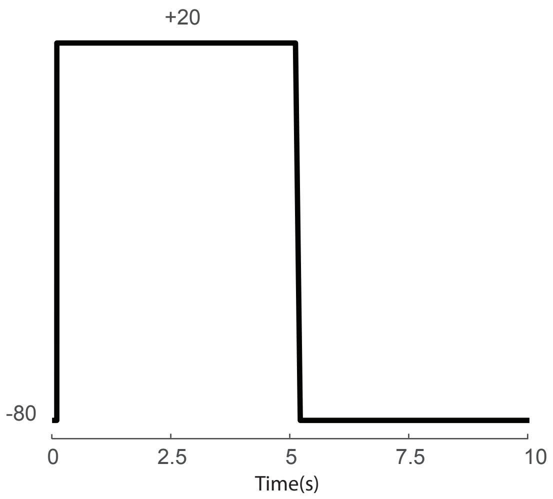


In Silico Kv11.1 Current

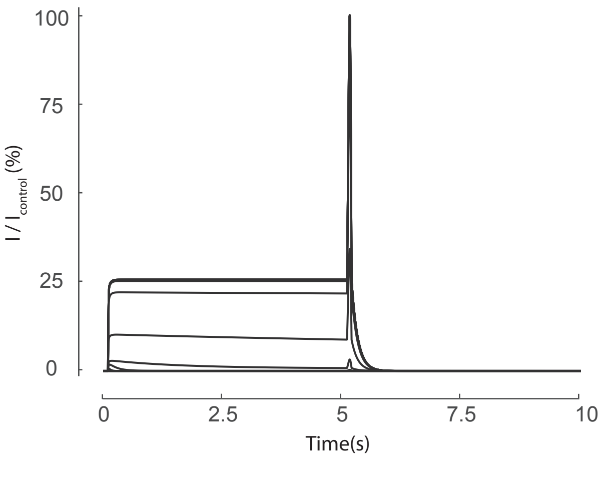

### Protocol (10)

Voltage Waveform

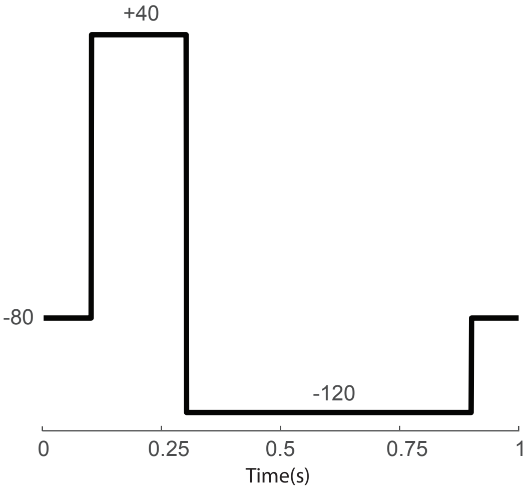


In Silico Kv11.1 Current

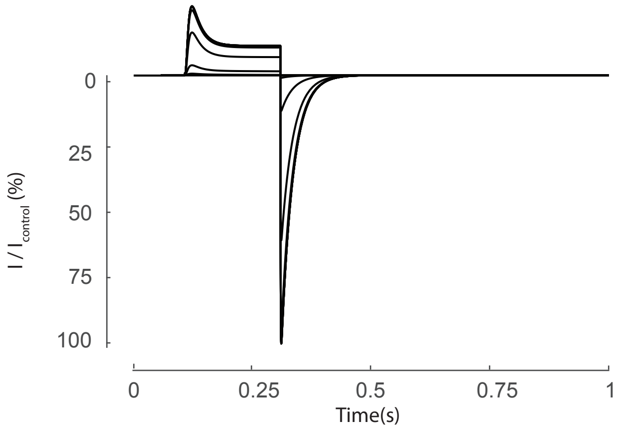


In Vitro Kv11.1 Current

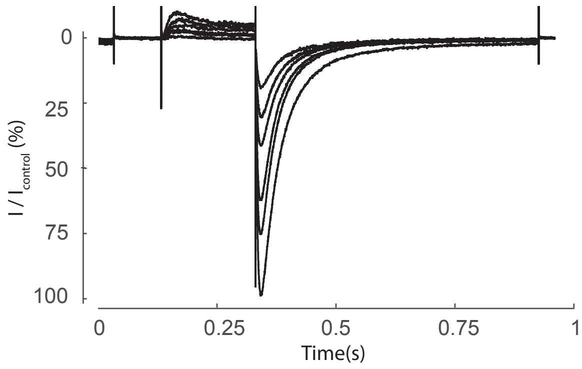

### Protocol (11)

Voltage Waveform

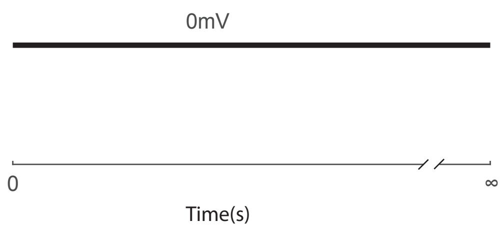


In Silico Kv11.1 Current

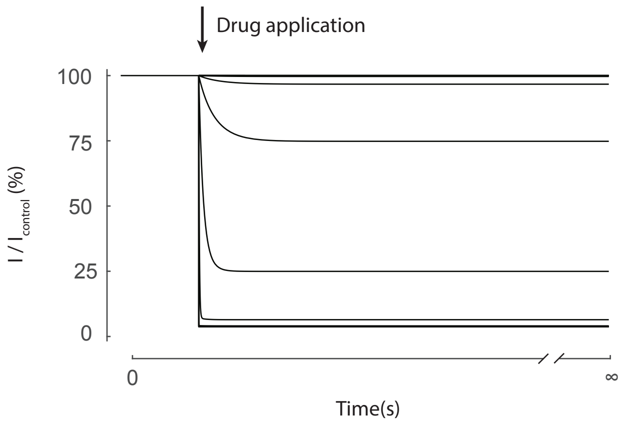


In Vitro Kv11.1 Current

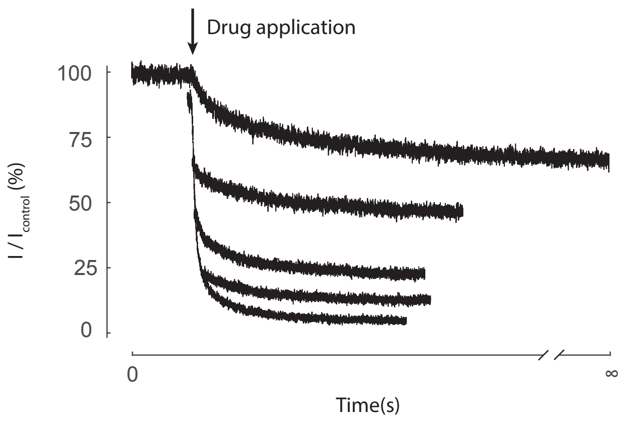

### Protocol (12)

Voltage Waveform

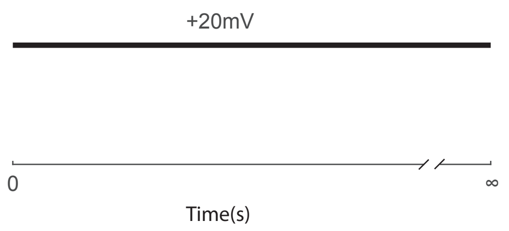


In Silico Kv11.1 Current

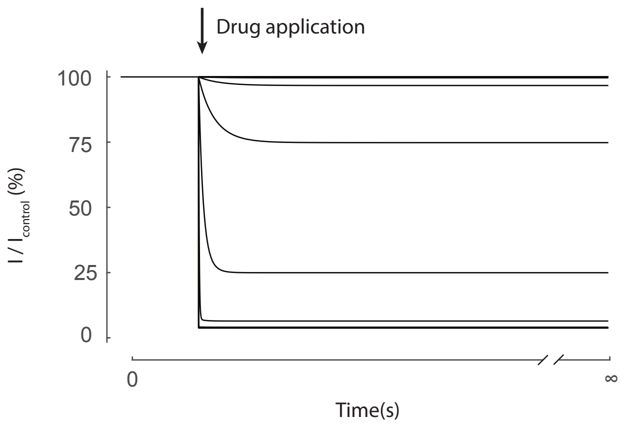


In Vitro Kv11.1 Current

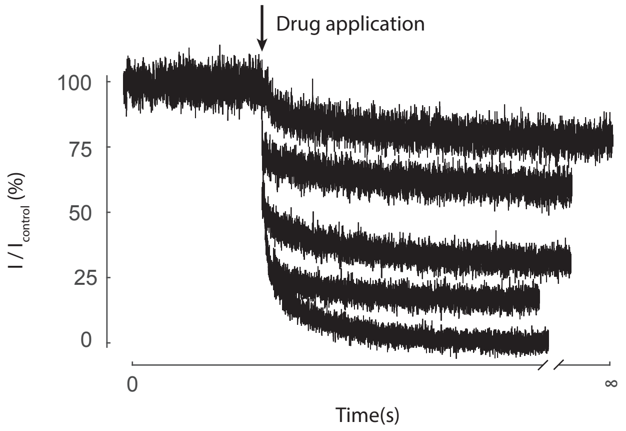

### Protocol (13)

Voltage Waveform

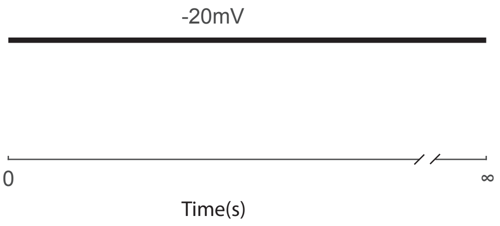


In Silico Kv11.1 Current

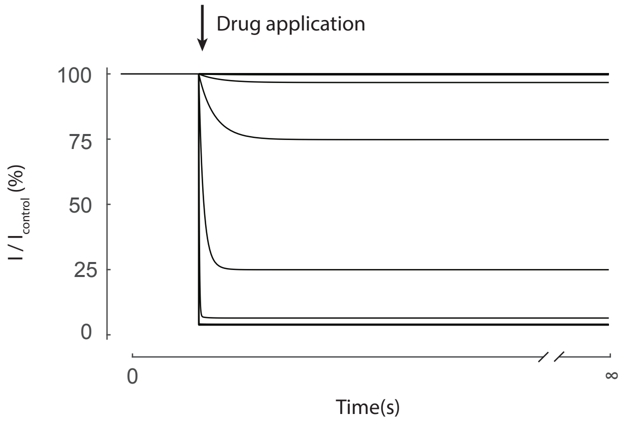


In Vitro Kv11.1 Current

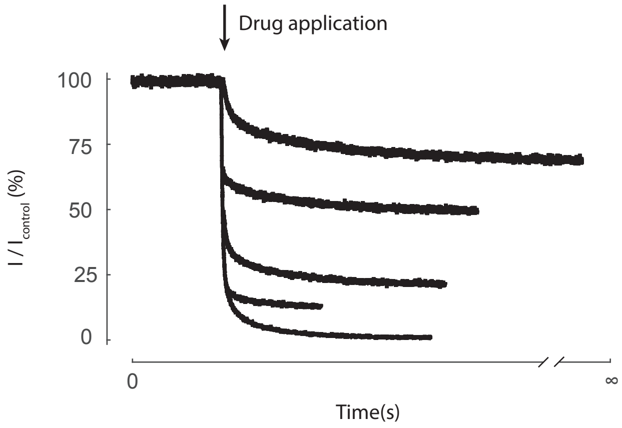

**Table 1**

| | Protocol | | | | | | | | | | |
|---|---|---|---|---|---|---|---|---|---|---|---|
| | (8) | (12) | (13) | (1) | (1)* | (10) | (10)* | (11) | (11)* | $K_{O/I}$ | $K_{O/I}$* |
| **verapamil** | 690.7 | 389.3 | 434 | 379.7 | 452.51 | 1112 | 262.14 | 537.1 | 397.11 | ~ | 1.97 |
| 95% CI | 348.7 - 1329 | 332.1 - 454.8 | 365.2 - 512.8 | 267.5 to 531 | | 899.5 - 1371 | | 444.5 - 644 | | | |
| Hill coefficient | -0.55 | -1.09 | -1.05 | -1.08 | | -1.07 | | -1.02 | | | |
| **cisapride** | 144.9 | 17.09 | 15.23 | 24.66 | 23.00 | 95.3 | 4.02 | 16.76 | 15.55 | ~ | 0.26 |
| 95%CI | 74.7-215.2 | 13.26 - 21.28 | 12.48 - 18.18 | 21.66 to 28.08 | | 78.93 - 115.3 | | 12.62 - 21.27 | | | |
| Hill coefficient | -0.73 | -0.92 | -1.23 | -1.13 | | -0.92 | | -0.90 | | | |
| **clozapine** | 1528 | 1299 | 1620 | 1429 | 1482.37 | 2600 | 1797.28 | 1654 | 1332.68 | 1.12 | 1.20 |
| 95%CI | 1180 - 1937 | 1096 - 1526 | 1354 - 1925 | 1318 to 1547 | | 2452 - 2758 | | 1317 - 2051 | | | |
| Hill coefficient | -0.81 | -0.96 | -0.97 | -1.04 | | -0.86 | | -0.93 | | | |
| **terfenadine** | 152.9 | 165.3 | 62.38 | 54.68 | 130.38 | 82.74 | 22.17 | 114.1 | 98.98 | ~ | 69.67 |
| 95% CI | 127.1 - 184.8 | 130.2 - 214 | 54.8 - 70.9 | 47.71 to 62.69 | | 59.55 - 114.9 | | 88.39 - 155.8 | | | |
| Hill coefficient | -0.95 | -1.23 | -1.67 | -1.39 | | -0.86 | | -1.38 | | | |

**Supplementary Table 1.** In vitro measured IC50s, their 95% confidence intervals and Hill coefficients for the four drugs (verapamil, cisapride, clozapine & terfenadine) are shown for the training protocols (8), (12) & (13) and the test protocols (1), (10) & (11). The test protocols are paired with their *in silico* IC50 prediction (denoted by *). KO/I and the *in silico* prediction KO/I* are also shown for each of the four drugs. IC50 values are show in nM.

Table 2

#### Verapamil

| 1st Author | year | $IC_{50}$ (nM) | cell line | $V_h$ | $V_1$ (mV) | $V_1$ (s) | $V_2$ (mV) | $V_2$ (s) | CL | [K]o (mM) | Temp (°C) |
|---|---|---|---|---|---|---|---|---|---|---|---|
| Chouabe | 1998 | **830** | COS-7 | -80 | +50 | 2 | -40 | 2 | 10 | 5 | 22 |
| Zhang | 1999 | **143** | HEK293 | -80 | +20 | 4 | -50 | 6 | 15 | 4 | 22 |
| Kutchinsky | 2003 | **450** | CHO | -90 | +20 | 2 | -50 | 2 | 10 | 4 | |
| Ridley | 2004 | **215.4** | HEK293 | -80 | 0 | 10 | | | | 4 | 37 |
| Wible | 2005 | **136** | HEK293 | -80 | +20 | 2 | -50 | 2 | 10 | | 22 |
| Okada | 2015 | **201.3** | CHO | -80 | +20 | 2 | -50 | 2 | 15 | 4 | 22 |
| Crumb | 2016 | **499** | CHO | Ventricular AP | | | | | 10 | 1 | 36 |
| Windley | 2017 | **399.5** | CHO | -80 | 0 | 10 | | | | 5 | 22 |

**Table 2.** Previously published $IC_{50}$ values for verapamil

**Table 3**

#### Cisapride

| 1st Author | year | $IC_{50}$ (nM) | cell line | $V_h$ | $V_1$ (mV) | $V_1$ (s) | $V_2$ (mV) | $V_2$ (s) | CL | [K]o (mM) | Temp (°C) |
|---|---|---|---|---|---|---|---|---|---|---|---|
| Rampe | 1997 | **45** | Mouse L cell | -80 | +20 | 2 | -40 | 1.6 | | 5 | 22 |
| Rampe | 1997 | **7** | Mouse L cell | -80 | +20 | 20 | | | | 20 | 22 |
| Mohammad | 1997 | **7** | HEK293 | -80 | +10 | 10 | -50 | 5 | 30 | 4 | 22 |
| Walker | 1999 | **16** | CHO | -75 | +25 | 3.9 | -55 | 5 | 10 | 4.8 | 22 |
| Walker | 1999 | **24** | CHO | -75 | +25 | 3.9 | -55 | 5 | 10 | 4.8 | 22 |
| Potet | 2001 | **240** | COS-7 | -80 | 10 | 0.5 | -60 | 0.5 | 3 | 4 | 35 |
| Wang | 2003 | **14** | HEK293 | -80 | +40 | 1 | -50 | | | 4 | 37 |
| Kirsch | 2004 | **26** | HEK293 | -80 | +20 | 2 | -50 | 2 | 10 | 4 | 22 |
| Guth | 2004 | **5** | HEK293 | -80 | | | | | | | 22 |
| Martin | 2004 | **18** | HEK293 | -80 | 0 | 3 | -50 | 5 | 15 | 5 | 35 |
| Fossa | 2004 | **15** | HEK293 | -80 | +20 | 1 | ramp | 0.5 V/s | 4 | 4 | 37 |
| Furuta | 2004 | **30** | HEK293 | -70 | 0 | 0.75 | -50 | 0.75 | | 4 | 37 |
| Chiu | 2004 | **7** | HEK293 | -75 | 10 | 0.5 | -40 | 0.5 | 10 | 4 | 37 |
| Kirsch | 2004 | **23** | HEK293 | -80 | 20 | 2 | -50 | 2 | 10 | 4 | 37 |
| Kirsch | 2004 | **27** | HEK293 | -80 | 20 | 1 | ramp | 0.5 V/s | 5 | 4 | 37 |
| Rezazadeh | 2004 | **4** | HEK293 | -80 | +20 | 4 | -50 | 3 | 11 | 5 | |
| Wible | 2005 | **27** | HEK293 | -80 | +20 | 2 | -50 | 2 | 10 | | 22 |
| Lin | 2005 | **8** | HEK293 | -80 | +50 | 4 | -50 | 5 | 15 | 0 | 23 |
| Lin | 2005 | **24** | HEK293 | -80 | +50 | 4 | -50 | 5 | 15 | 5 | 23 |
| Lin | 2005 | **109** | HEK293 | -80 | +50 | 4 | -50 | 5 | 15 | 135 Cs | 23 |
| Lin | 2005 | **174** | HEK293 | -80 | +50 | 4 | -50 | 5 | 15 | 0Cs | 23 |
| Lin | 2005 | **192** | HEK293 | -80 | +50 | 4 | -50 | 5 | 15 | 5 Cs | 23 |
| Lin | 2005 | **196** | HEK293 | -80 | +50 | 4 | -50 | 5 | 15 | 135 Cs | 23 |
| Toga | 2007 | **9** | HEK293 | -80 | +40 | 1 | ramp | 0.5 V/s | 4 | 4 | 23 |
| Perrin | 2008 | **21** | CHO | -80 | +20 | 3 | -110 | 0.5 | 10 | 5 | 22 |
| Okada | 2015 | **14.7** | CHO | -80 | +20 | 2 | -50 | 2 | 15 | 4 | 22 |
| Windley | 2016 | **23.1** | CHO | 0mV | | | | | | 5 | 22 |
| Crumb | 2016 | **12** | CHO | Ventricular AP | | | | | | 1 | 36 |
| Kang | 2016 | **15** | CHO | -80 | +20 | 2 | -40 | | 10 | 4 | 37 |
| Windley | 2017 | **18.9** | CHO | -80 | 0 | 10 | | | | 5 | 22 |

**Table 2.** Previously published $IC_{50}$ values for cisapride

### Table 4

#### Clozapine

| 1st Author | year | $IC_{50}$ (nM) | cell line | $V_h$ | $V_1$ (mV) | $V_1$ (s) | $V_2$ (mV) | $V_2$ (s) | CL | [K]o (mM) | Temp (°C) |
|---|---|---|---|---|---|---|---|---|---|---|---|
| Tie | 2000 | **2630** | CHO | -75 | +25 | 3.9 | -55 | 5 | 10 | 4.8 | 22 |
| Ekins | 2002 | **320** | HEK293 | -75 | +10 | 0.5 | -40 | 0.5 | 10 | 4 | 37 |
| Lee | 2009 | **2500** | HEK293 | -70 | +40 | 4 | -60 | | | 5.4 | 36 |
| Kramer | 2013 | **2300** | HEK293 | -80 | +40 | 2 | -40 | 2 | 10 | 4 | 22 |
| Hill | 2014 | **2800** | CHO | 0 | | | | | | | 22 |

**Table 4.** Previously published $IC_{50}$ values for clozapine

### Table 5

#### Terfenadine

| 1st Author | year | $IC_{50}$ (nM) | cell line | $V_h$ | $V_1$ (mV) | $V_1$ (s) | $V_2$ (mV) | $V_2$ (s) | CL | [K]o (mM) | Temp (°C) |
|---|---|---|---|---|---|---|---|---|---|---|---|
| Crumb | 2000 | **204** | HEK293 | -75 | +10 | 0.4 | -40 | 0.4 | 10 | | 36 |
| Wang | 2003 | **9** | HEK293 | -80 | +40 | | -50 | | | | 36 |
| Martin | 2004 | **16** | HEK293 | | | | | | 2 | 5 | 37 |
| Wible | 2005 | **8** | HEK293 | -80 | +20 | 2 | -50 | 2 | 10 | | 22 |
| Perrin | 2008 | **61.4** | CHO | -80 | +20 | 3 | -110 | 0.5 | 10 | 5 | 22 |
| Friemel | 2010 | **27.7** | HEK293 | -80 | +20 | 2 | -40 | 2 | 10 | 5.6 | 22 |
| Tanaka | 2014 | **30.6** | HEK293-hergGFP | -80 | +20 | 1.5 | -50 | 1.5 | 15 | 4 | 37 |
| Okada | 2015 | **98.5** | CHO | -80 | +20 | 2 | -50 | 2 | 15 | 4 | 22 |
| Badyra | 2015 | **70** | HEK293 | -70 | -50 | 0.1 | +40/-50 | 02-Feb | 10 | 2.5 | 37 |
| Crumb | 2016 | **19** | CHO | Ventricular AP | | | | | | 1 | 36 |
| Kang | 2016 | **14** | CHO | -80 | +20 | 2 | -40 | | 10 | 4 | 37 |
| Windley | 2017 | **13.1** | CHO | -80 | 0 | 10 | | | | 5 | 22 |

**Table 5.** Previously published $IC_{50}$ values for terfenadine
